# Programmed Delayed Splicing: A Mechanism for Timed Inflammatory Gene Expression

**DOI:** 10.1101/443796

**Authors:** Jacob S. Dearborn, Luke Frankiw, Damas W. Limoge, Christian H. Burns, Logan Vlach, Patricia Turpin, Tylar Kirch, Zachary D. Miller, William Dowell, Sylvester Languon, Yvette Garcia-Flores, Robert C. Cockrell, David Baltimore, Devdoot Majumdar

## Abstract

Inflammation involves timed gene expression, suggesting that the fine-tuned onset, amplitude, and termination of expression of hundreds of genes is of critical importance to organismal homeostasis. Recent study of post-transcriptional regulation of inflammatory gene expression led to the suggestion of a regulatory role for pre-mRNA splicing. Here, using a hybrid capture approach to purify incompletely spliced, chromatin-associated pre-mRNAs, we use deep sequencing to study pre-mRNA splicing of the NF-*𝜅*B transcriptome. By freezing transcription and examining subsequent splicing of complete transcripts, we find many introns splice tens to hundreds of times slower than average. Investigating the basis of these delays, we focused on evolutionarily conserved introns with suboptimal splice donor sequences and found that strengthening these donor sites by as few as two nucleotides in minigene reporter assays markedly increased gene expression for several targets. This suggests that such sites can act as timing elements that both delay mRNA production and limit expression amplitude. To broaden this mechanistic view, we applied deep learning sequence-to-function models with feature attribution to identify additional regulatory sequences—both intronic and exonic—that may contribute to delayed splicing through mechanisms independent of donor site strength. This integrated approach revealed non-canonical motifs enriched in slow-splicing introns, pointing to a broader repertoire of cis-elements that can fine-tune transcript maturation during inflammation. Together, these findings support a model in which the temporal regulation of pre-mRNA splicing serves as a layer of control in inflammatory gene expression, and raise the possibility that similar timing mechanisms operate in other rapid-response transcriptional programs.

## Introduction

Gene expression in response to an inflammatory stimulus begins rapidly and is tightly controlled by conventional means (transcription and protein turnover; (Chen and Chen, 2013; Gautier et al., 2012; Smale et al., 2014)) and by an expanding list of modalities that have gained in appreciation as being general regulatory strategies (RNA stabilization, RNA deadenylation, ribosomal regulation, microRNA regulation, as examples; (Hao and Baltimore, 2009; Leppek et al., 2013; O’Connell et al., 2012; Wan et al., 2007)). We and others have recently investigated the role of RNA splicing kinetics—independently of alternative splicing—in gene expression (Hao and Baltimore, 2013; Pandya-Jones et al., 2013; Rabani et al., 2011, 2014). In macrophages, an inflammatory stimulus leads to upregulation of expression of pre-mRNAs from hundreds of genes, providing an experimentally favorable system to investigate whether differential kinetics of pre-mRNA splicing may control the timing of gene expression following an inducing stimulus.

Pre-mRNA conversion to mRNA has been implicated in regulation of gene expression in diverse systems. As part of the cellular response to various environmental stressors, mRNAs for ribosomal proteins were shown to be downregulated due to decreased splicing efficiency in yeast (Bergkessel et al., 2011). Global changes in efficiency of pre-mRNA splicing have been shown to be a developmental prerequisite for *Drosophila* early embryonic development (Guilgur et al., 2014). The developing vertebrate embryo obeys a “segmentation clock” determining body segment length whose very timing relies on delays attributable to control of the splicing rate of the *Hes7* transcriptional repressor (Takashima et al., 2011).

In certain well-studied cases, as with the cytokine TNF*𝛼*, regulatory mechanisms modulating RNA levels exert significant physiological effects (Eissa et al., 1996; Hargreaves et al., 2009). The insight that TNF*𝛼* contains AU-rich elements in its 3^′^ untranslated region that act as mRNA degradation signals (Han et al., 1990), and subsequent observations that a mouse in which these AU-rich elements were removed resulted in a robust autoimmune phenotype (Kontoyiannis et al., 1999), was an early indication of the importance of precisely tuned mRNA levels in the regulation of inflammation to avoid autoimmunity. Given the role of pre-mRNA splicing in biogenesis of mature mRNA, we and others (Hao and Baltimore, 2013; Bhatt et al., 2012; Cho et al., 2014; Davis-Turak et al., 2015; Grabherr et al., 2011) suggested that regulation of splicing kinetics may influence the gene expression kinetics that define the inflammatory cascade. Consistent with this idea, Braunschweig et al. (2014) demonstrated that intron retention is a widespread and regulated feature of mammalian transcriptomes, often associated with reduced mRNA abundance and subject to tissue-specific control.

Deep learning models have rapidly advanced our ability to interpret noncoding regions of the genome by learning complex regulatory logic directly from DNA sequence. Early models such as DeepSEA demonstrated that convolutional neural networks could predict the effects of noncoding variants on transcription factor binding and chromatin state, laying a foundation for sequence-to-function inference (Zhou and Troyanskaya, 2015). Building on this, SpliceAI adapted deep convolutional architectures to model splicing dynamics directly from primary sequence, achieving high accuracy in predicting splice site usage and the effects of sequence variants on splicing (Jaganathan et al., 2019). More recent models such as Enformer and Borzoi have expanded this paradigm further by predicting RNA-based outputs, including gene expression levels, transcript isoform usage, and transcription start site precision (Avsec et al., 2021; Linder et al., 2025). These models also support feature attribution through interpretable machine learning approaches such as saliency mapping and deepSHAP, allowing inference of sequence elements—such as transcription factor binding sites and splicing regulators—that most influence RNA output. Together, these approaches provide an unprecedented modeling context—spanning hundreds of kilobases—that enables the systematic dissection of sequence-level determinants of intron retention and the broader landscape of RNA processing.

To examine the timing of intron removal from 230 different transcripts induced by TNF*𝛼*in macrophages, we have developed a method for highly enriching transcript populations for mRNAs of interest, which is followed by deep sequencing of the largely pre-mRNA populations we purify. The induction of transcripts and removal of introns can be quantified with precision, with lifetimes of introns determined by blocking transcription early after induction. Among genes whose mRNAs appear more slowly after induction, we identify ones containing introns with poor binding sites for splicing factor U1, usually finding one per transcript. We call these “bottleneck” introns. Among the most rapidly induced genes we find no such introns. We show that in these pre-mRNAs, the sequence of the U1 binding site is critical for the speed of intron removal by building mini-genes with “repaired” introns and showing that these splice at the canonical rate (identified as about 20 seconds after polymerase has passed that point). To complement these findings and explore regulatory determinants at larger scale, we apply a deep learning model trained on genomic sequence to predict splicing dynamics. Feature attribution methods identified sequence features associated with the splicing delays. We observe and model fine-tuning improved detection of patterns associated with intron retention. We propose that bottleneck introns are important for determining either the rate of degradation of pre-mRNAs or the rate of appearance of mature mRNA or both.

## Results

### Hybrid Capture of Chromatin-Associated Transcripts

It has been established that pre-mRNA is highly enriched in the chromatin-associated, polyadenylated RNA fraction (Tilgner et al., 2012). To examine splicing events in pre-mRNAs, we performed time-course experiments with TNF-stimulated bone marrow-derived macrophages (BMDMs), isolating RNA after biochemical separation of chromatin-associated material (Bhatt et al., 2012). Using a hybrid-capture approach, we targeted sequencing toward transcripts of 230 genes previously identified as TNF-induced inflammatory mRNAs (Ramirez-Carrozzi et al., 2009).

The hybrid capture strategy, based on a published approach (Engreitz et al., 2013), involved: (1) microarray printing of 12,000 150-bp ssDNAs designed from tiled fragments of the last exon of each gene of interest; (2) conversion to a pool of biotinylated ssRNA probes using PCR followed by in vitro transcription with biotinylated ribonucleotides; (3) hybridization of ssRNA pools to cDNA from each biological experiment; and (4) streptavidin-coated bead-mediated capture of the transcripts of interest (Fig. S1).

This approach resulted in a ∼30-fold enrichment of genes of interest, with 70% of the sequenced reads corresponding to the genes of interest; RNA submitted only to poly(A) selection contained only 2% of such reads (Fig. S2A).In this way we could analyze 1098 introns (after TPM filter >100) from genes induced by an inflammatory stimulus (Fig. S2B). The selected chromatin-associated transcripts collected across time points after induction are shown as read-density histograms, displaying sequencing read abundance along each gene to reveal exon–intron structure. In Fig. 1A, *Nfkbia* (encoding I*𝜅*B*𝛼*) is shown over 1 hour after TNF induction, with log_10_-scaled densities normalized within each time point to highlight transcriptional kinetics. New transcription becomes evident by 6 minutes, especially on an unnormalized linear scale (Fig. S3). The corresponding logscaled histograms permit visualization of the intronic signal as a function of time after induction.

**Figure 1.**
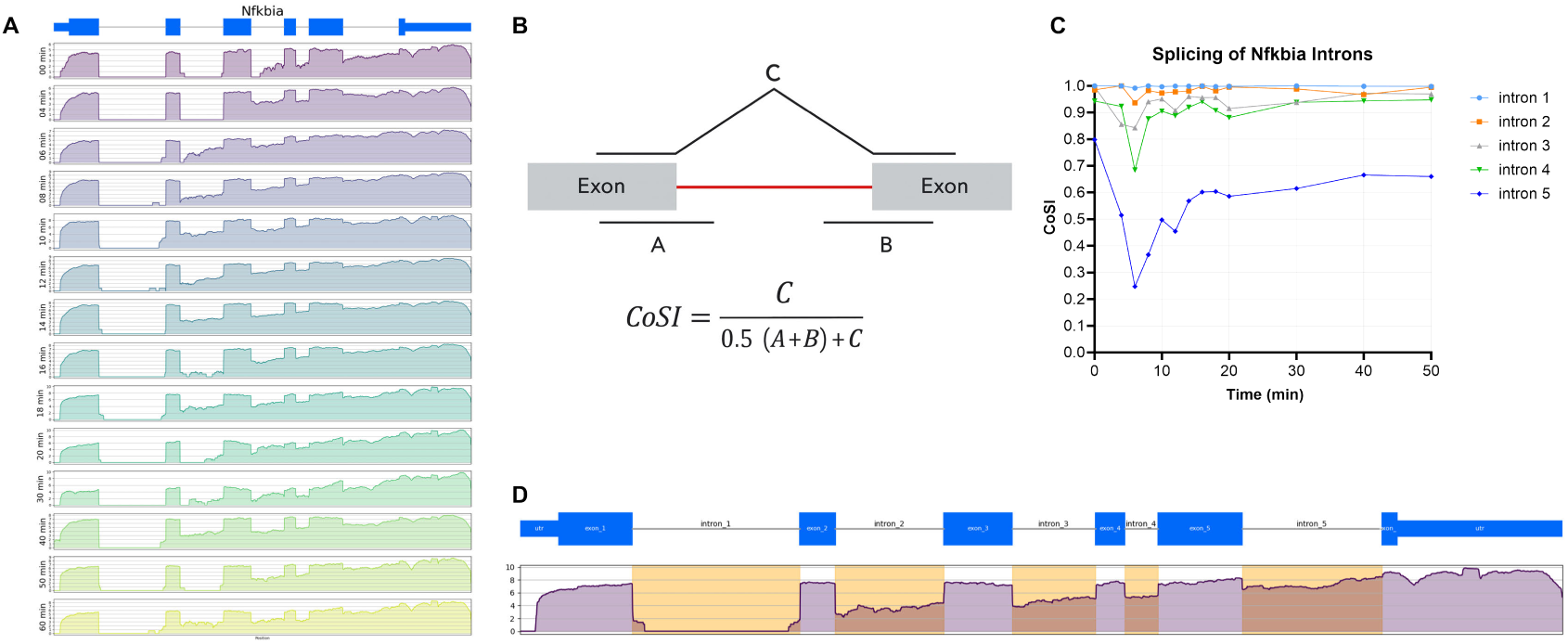
Sequencing of complete, chromatin-associated pre-mRNA during inflammatory stimulus reveals differential splicing dynamics among introns of IKB*𝛼*. (A) Histogram of reads corresponding to the TNF-induced expression and splicing of IKB*𝛼* pre-mRNA of BMDMs. RNA-seq was performed on chromatin-associated RNA, enriched for NF*𝜅*B genes as a function of a TNF stimulation time course, time shown in minutes after stimulation. Reads are histogrammed in log_10_ scale and normalized to each time point’s maximum value. (B) The Coefficient of Splicing (CoSI) metric quantifies extent of splicing as a function of time, expressed as a ratio of reads from each splice junction to total junctional reads. Dynamics of IKB*𝛼* splicing as a function of each intron’s CoSI is shown (C), where 1 = spliced and 0 = unspliced, with corresponding introns highlighted in a sample time point. (D) Differential dynamics of splicing for each *Nfkbia* intron are further demonstrated in the coverage plot for the transcript.

Furthermore, from Fig. 1A it is evident that at all time points following induction, the 5^′^-proximal introns have been totally removed from the sequenced transcripts, indicating that the selection against partial transcripts is quite complete. Whereas excision of the first intron is always observed, the middle three introns are seen at intermediate states of excision at all time points such that intron definition for these introns is readily observable from read density histograms. We attribute this first exon excision largely to co-transcriptional splicing, consistent with other genome-wide splicing studies (Bentley, 2014). Strikingly, the final intron deviates significantly in its kinetic trajectory, as its read density does not obey a similar relative reduction. This might be due to a lag in terminal intron splicing (Carrillo Oesterreich et al., 2010) or a feature of splicing that accompanies transcript release from chromatin.

### Quantifying Splice Completion Across the Transcriptome

To better quantify the observed dynamics, we adapted the Coefficient of Splicing (CoSI) (Fig. 1B), which quantifies the extent of splicing as a ratio of spliced to total (spliced and unspliced) junction reads such that CoSI values of ∼1 and ∼0 imply near-complete splicing and virtually unspliced states, respectively (Tilgner et al., 2012). Though we observed a decrease in read density as a function of distance from the 3^′^ end of the gene (Fig. S3), presumably as a consequence of premature termination of the reverse transcriptase during copying of the pre-mRNA, the use of CoSI allows for an intron-specific splicing score regardless of read densities at neighboring introns. Using the CoSI metric, a time course plotting of the extent of splicing showed a dip in CoSI at ∼6 minutes (Fig. 1C) corresponding to the aforementioned accumulation of new, unspliced transcripts. The splicing dynamics of each *Nfkbia* intron can be inferred from the CoSI dynamics, and the notable difference in splicing between the 5^′^ and 3^′^ introns (Fig. 1D) is demonstrated by the amplitude of the “dip” at 6 minutes and the time required for each intron to return to CoSI of 1.

A surprising heterogeneity in CoSI was observed among all inflammatory introns (Fig. 2A), implying diversity in their propensity to be spliced (Fig. 2). When considering all 1,024 introns in the chromatin-associated TNF time course we find most introns very near to CoSI ∼1 relatively soon after induction, indicating that most introns do not remain unspliced long after induction begins between 4–8 minutes (Fig. 2A). The observation that many TNF-responsive transcripts initiate within 4–8 minutes is consistent with that of other related studies (Bhatt et al., 2012). Unexpectedly, although the median CoSI value remains high, we identified considerable heterogeneity among introns, some at and remaining near CoSI values < 0.5, indicating relatively poor splicing, found very late into the time course.

**Figure 2.**
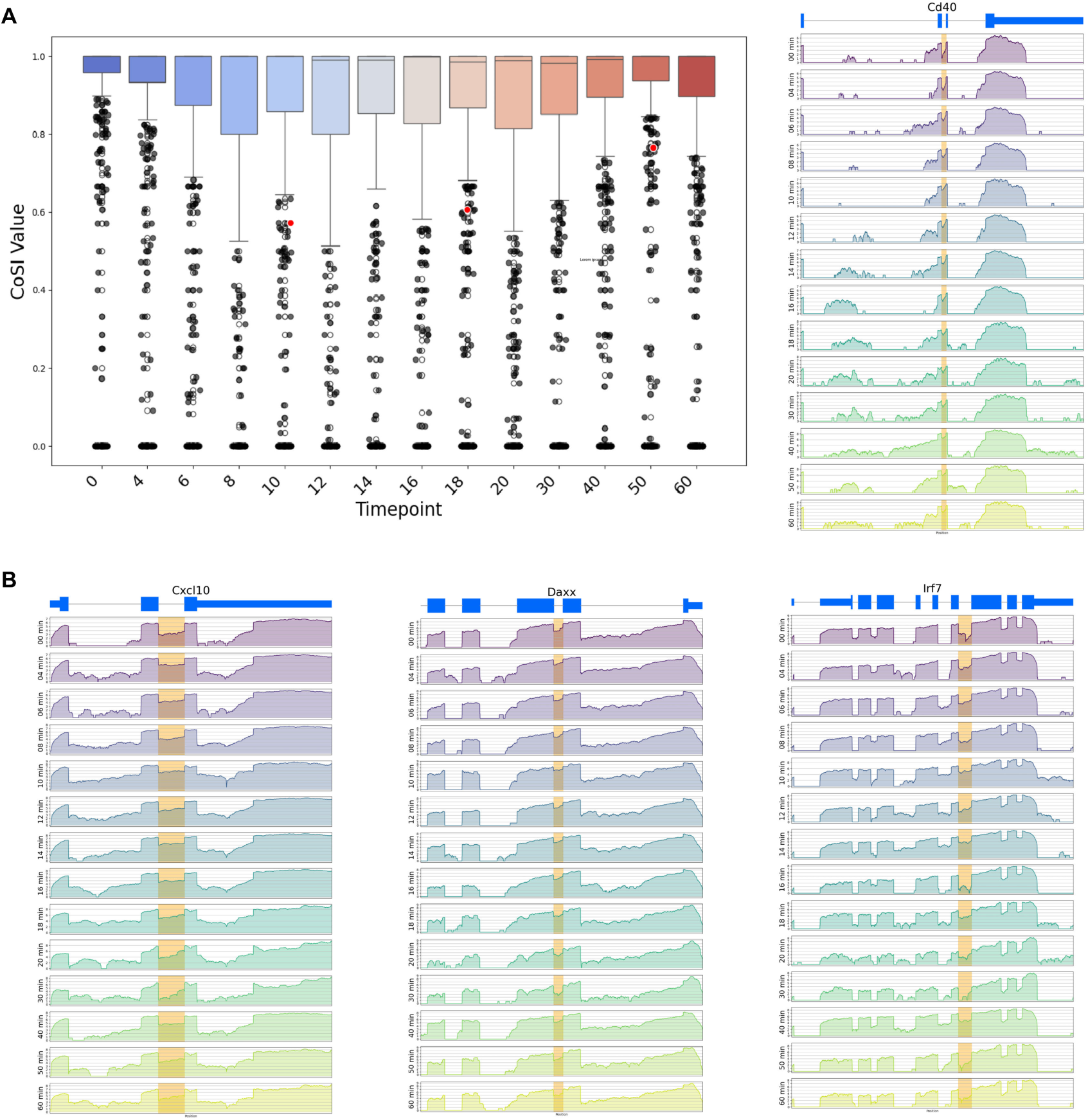
Heterogeneity of splicing at each intron reveals splicing “bottlenecks”. The CoSI of each intron per time point is shown as a function of the entire inflammatory mRNA dataset as a box–whisker plot (A). Each point represents an intron of one of 230 genes, revealing high rates of splicing (median CoSI indicated by bar near 1.0 for each time point) for most genes with significant outliers. As an example, *Cxcl10* intron 2 (red arrowhead) is represented by the datapoint with arrowhead, and a histogram of reads is shown to demonstrate the relatively unspliced nature of this intron, which is not involved in alternative splicing. (B) Several similar introns that are relatively unspliced are found throughout the inflammatory transcriptome; shown are bottleneck introns within *Cd40*, *Daxx*, and *Irf7* as examples in the context of their neighboring introns.

As an example of a poorly spliced intron, chemokine *Cxcl10* intron 2 (Fig. 2A) is notable as it remains poorly spliced despite clear excision of neighboring introns, remaining quite unspliced even ∼30 minutes after induction. It is possible that this intron undergoes splicing after its nascent chromatin-associated state, as is likely the case for the 3^′^-terminal intron of *Nfkbia*. It is also possible that this intron targets *Cxcl10* transcripts for degradation and the relatively fixed nature of intron 2’s splicing status throughout the time course is a function of a constant rate of degradation. Introns with low CoSI at late time points post-TNF induction were considered putative “bottleneck introns”—borrowing from the language that accompanied the discovery and characterization of slowly splicing U12-type introns (Patel et al., 2002). These introns were so slow to splice that they may intrinsically delay gene expression. Notably, the distribution of CoSI values of the entire dataset (Fig. 2A) was very broad, and though most introns spliced immediately, many introns showed evidence of splicing bottlenecks, noticeable by their significant deviation from mean CoSI. At 10 and 60 minutes post-induction, 14% and 11% of introns, respectively, had CoSI values below one standard deviation of the mean (0.86±0.25 and 0.91±0.19, respectively). Shown as examples are *Cd40*, *Cxcl10*, *Daxx*, and *Irf7* (Fig. 2A–B), genes whose immunological and inflammatory importance is well-established in studies with knockout mice (Honda et al., 2005; Lei et al., 1998; Michaelson et al., 1999).

### Measurements of Intron Splicing Half-Lives

To quantify splicing kinetics, we used Actinomycin D (actD) to freeze transcription and followed the loss of intron and accumulation of splice junctions. In these experiments, splicing was analyzed at many time points immediately following actD treatment on the same 230 transcripts of interest and selected using hybrid capture from the total pool of cellular RNA rather than chromatin-associated RNA. Fitting the accumulation of spliced transcripts (as measured by CoSI) with an exponential distribution, we were able to extrapolate intron excision half-lives (Fig. 3). Because total cellular RNA was used, observed rates were independent of chromatin localization. We find intron half-lives that range from 20–40 s (56% of introns splicing in this timing window) to several minutes, reflecting the considerable heterogeneity that is observed from CoSI differences.

**Figure 3.**
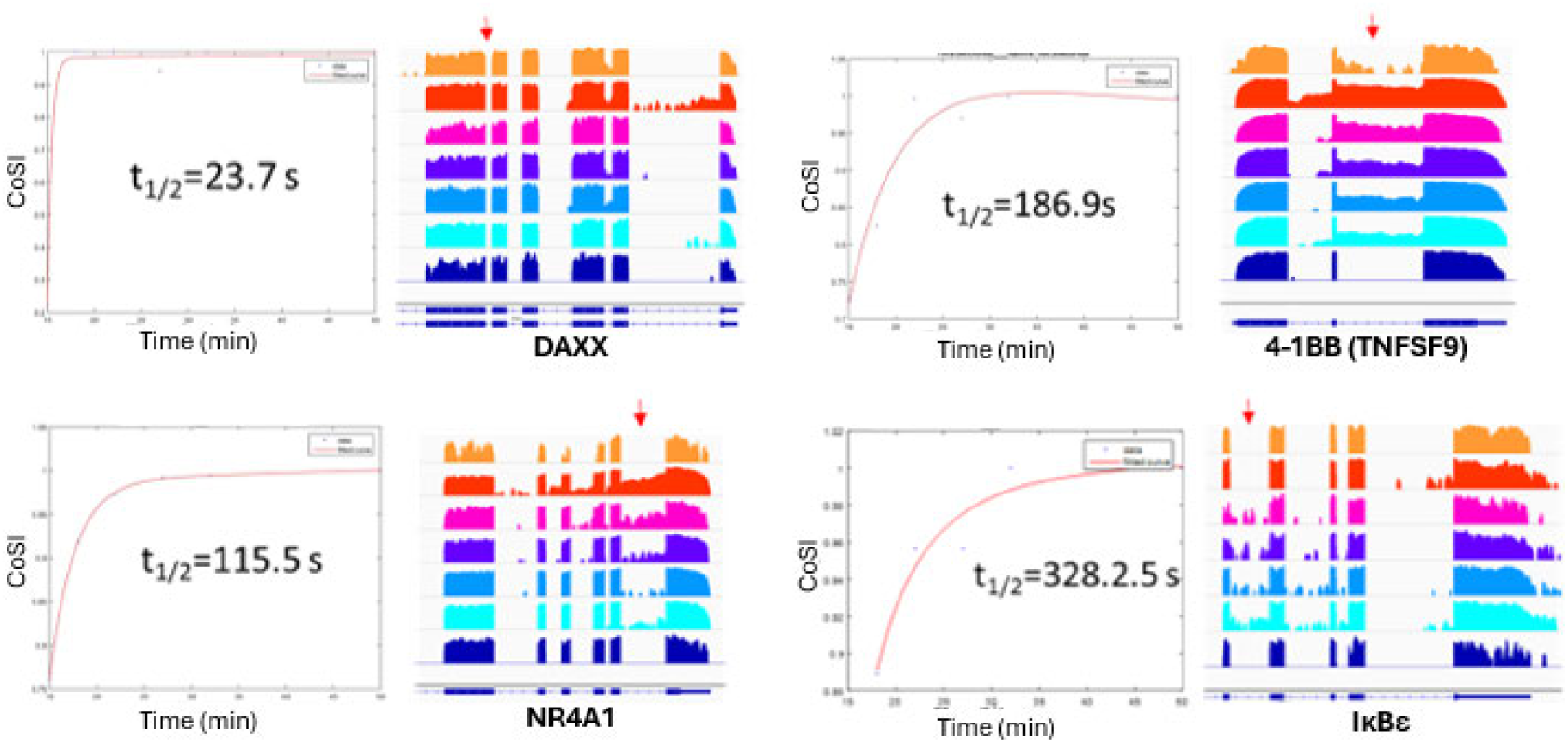
Splicing kinetics of inflammatory introns are heterogeneous, ranging from seconds to minutes. CoSI of introns representing various splicing rates are measured and fit to half-lives. Cells were treated with Actinomycin D, from which hybrid capture of genes of interest and sequencing was performed on total (unfractionated) RNA. Shown are four representative samples of splicing kinetics.

### A Multifaceted Approach for Identification of Determinants of Slow Splicing

Building on our observation that certain introns exhibit markedly delayed splicing by both CoSI dynamics and direct half-life measurements, we sought to uncover the underlying sequence features responsible for these delays. To this end, we employed a multifaceted strategy: (1) computational analysis of 5^′^ splice donor strength using MaxEntScan, (2) experimental testing of splicing efficiency with a minigene reporter assay, and (3) interpretable deep learning to identify additional cis-regulatory elements—beyond canonical splice sites—that may modulate splicing kinetics.

### Computational Analysis of Splice Donor Strength

A delay in splicing at certain sites could simply confer a delay in gene expression (by ∼5 minutes in the slow case shown in Fig. 3, I*𝜅*B*𝜖*), or, as is seen in yeast studies, it could result in both gene expression delay and gene expression diminution due to degradation of slowly splicing pre-mRNA (Koodathingal et al., 2010). Prior studies placed I*𝜅*B*𝜖* in a delayed splicing category (Hao and Baltimore, 2013), suggesting a pronounced splicing delay relative to rapidly induced genes. To understand a potential mechanistic basis for these differences in splicing time, each intron within our dataset was assessed computationally for the concurrence of its 5^′^ splice donor sequence to a consensus sequence (Pessa et al., 2006). The 5^′^ splice donor is a highly conserved sequence that directly base pairs with splicing factor U1 (Freund et al., 2005); deviation from the consensus sequence confers a significantly reduced ability to engage the splicing machinery.

A maximum entropy model (Yeo and Burge, 2004) was used to calculate an intron quality score measuring extent of deviation from the consensus splice sequence (Fig. S4). Among the inflammatory transcripts studied, many examples of introns with poor 5^′^ donor scores were identified such as *Irf7* and *Il12b*, where lower scores indicate significant deviation from consensus. We suggest that having non-consensus splice sites may be a regulatory mechanism affecting gene expression. We considered that splicing might show profound differences in the previously defined categories of induction (immediate/early/intermediate/late) characteristic of the inflammatory gene expression kinetics (Bhatt et al., 2012). We found that the “immediate” genes showed consistently fast splicing (highest CoSI values) at all of their introns except the most 3^′^ intron, whereas the other three groups shared similar CoSI distributions (Fig. S6). Using the bioinformatics “intron quality score” we also found that the introns of the later gene classes have significantly lower scoring 5^′^ splice donor sequences (Fig. S4). Therefore, from experimental measurement of splicing and sequence-based prediction, genes expressed immediately following inflammatory stimulus are spliced faster, whereas all other inflammatory genes have a complex and heterogeneous distribution of splicing efficiency that does not stratify cleanly into the later kinetic categories (early/intermediate/late). Slowly splicing introns are found throughout these later kinetic categories in similar abundance, perhaps playing very gene-specific roles in diverse kinetic categories.

### Experimental Validation with Minigene Fluorescent Reporter Assay

To test whether delays in splicing result in changes to gene expression, we identified a set of introns with the following criteria: (1) introns that splice poorly as defined by RNA-Seq, (2) introns that contain a low-scoring (non-consensus) 5^′^ splice donor, and (3) introns whose weak 5^′^ splice donor is evolutionarily conserved across many mammalian species. These introns were tested in the context of a splicing reporter expressed on a bidirectional promoter (Mukherji et al., 2011). For each intron of interest, the reporter construct consists of a single transcript containing: (1) the 5^′^ neighboring exon from the gene of interest, the intron of interest, and the 3^′^ neighboring exon; (2) a 2A “self-cleaving” peptide; and (3) the GFP gene. In the opposite orientation but from the same promoter, a blue fluorescent protein (BFP) mRNA is made in equal amounts to the intron-GFP construct. GFP fluorescence of cells transfected with this reporter is a readout of splicing efficiency of the intron-GFP construct, when normalized to BFP fluorescence levels. Transfected into HEK293T cells and expressed for 24 hours, this bidirectional reporter enabled us to understand, at steady state, whether gene expression is affected by slow splicing introns. Measured using flow cytometry, the slope of the line corresponding to BFP:GFP ratios provides a relative metric of splicing efficiency, where slopes ∼0.9 and ∼0.1 imply efficient and inefficient splicing, respectively.

To test the effect of a poor splice donor, we “rescued” some poorly splicing introns in the context of the reporter by mutation to consensus splice donor “GTAAG.” For instance, for *IL12* intron 3, the splice donor sequence of “GTAAT” that is conserved among many mammalian species was altered to “GTAAG” (Fig. S5). Expression levels of the reporter construct with the wild-type *IL12* intron were found to be about half (57%) of the levels of the same construct with a single base pair alteration to make the stronger splice donor. *IRF7* intron 5 was tested against a “fixed” intron as well as a wild-type intron from an actin gene, both resulting in two-fold improvements of gene expression. In one case, expression of TFEC (transcription factor EC) was not altered by splice site repair, suggesting that other mechanisms beyond 5^′^–splice site deficiencies may be involved in mediating slow splicing. Generally, when the BFP:GFP slopes of wild-type and mutated introns were compared by taking a ratio of their slopes, a change of ∼2 nucleotides dramatically altered the slope of the line (Fig. 4).

**Figure 4.**
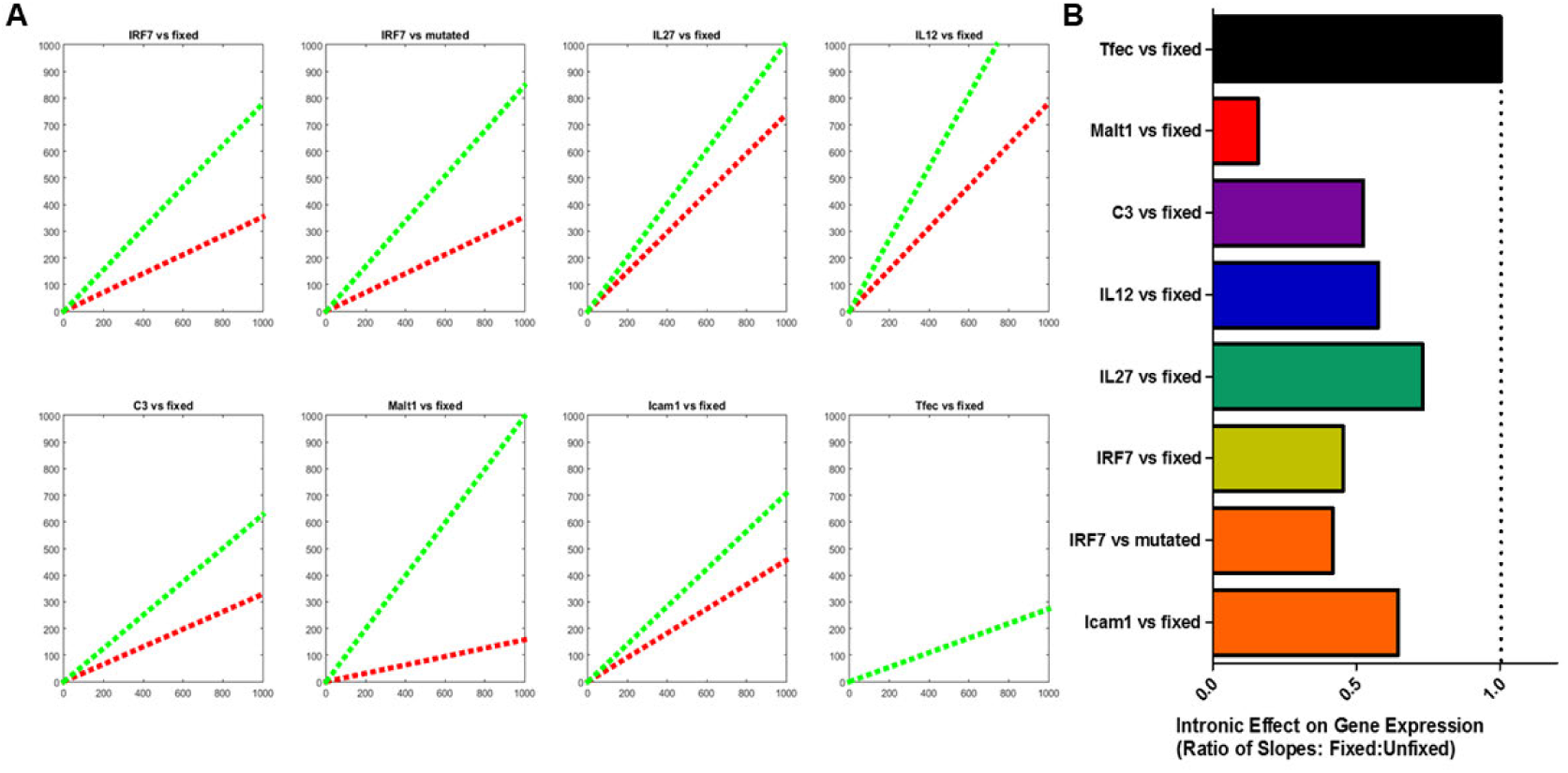
Bottleneck introns can be repaired and account for significant alterations to gene expression. (A) Intron–GFP splicing reporters for each wild-type intron (red) and modified intron (green) are shown as BFP:GFP ratio. (B) Ratio of WT:fixed slopes is shown; whereas *Tfec* expression is not altered by improved 5^′^ sequence, *Malt1* intron sequence is significantly impaired owing to its 5^′^ donor sequence, exhibiting a roughly five-fold impairment in gene expression due to the 5^′^ splice donor.

### Deep Learning–Based Discovery of Non-Canonical Regulatory Motifs

To extend our understanding of regulatory features beyond canonical splice donor sequences, we applied a supervised regulatory sequence model (Borzoi) to investigate sequence determinants associated with slow splicing. Borzoi is a model originally trained on steady-state RNA profiles across diverse cell types and tissues, which may limit its ability to capture stimulus-specific regulatory dynamics at play during TNF*𝛼*-induced intron retention. Fine-tuning of this model offers a way to adapt a large, pre-trained sequence model to a specialized context by continuing training on domain-specific datasets, leveraging the general regulatory knowledge already encoded in the model weights while learning new patterns relevant to the task at hand. Fine-tuning has been shown to enhance Borzoi’s performance in other specialized contexts, including tissue-specific expression, transcription factor knockdowns, and cellular aging (Yuan et al., 2025). We fine-tuned it using RNA-seq coverage data from BMDMs at 18, 20, and 30 minutes post-TNF*𝛼*induction—timepoints that capture dynamic intron retention. Fine-tuning was performed over 20 epochs using the Adam optimizer (learning rate = 1 × 10^−6^, MSE loss). To evaluate generalization, chromosomes 10 and 11 were held out for training and used for validation and testing, respectively. Model loss steadily decreased over the training period for both datasets (Fig. 5A), and Pearson correlation (reflecting the strength of their linear relationship) between predicted and observed expression improved from *𝑟*= 0.51 (*𝑝* = 4.44 × 10^−16^) for the pretrained model to *𝑟*= 0.61 (*𝑝* = 4.84 × 10^−24^) after fine-tuning (Fig. 5B).

**Figure 5.**
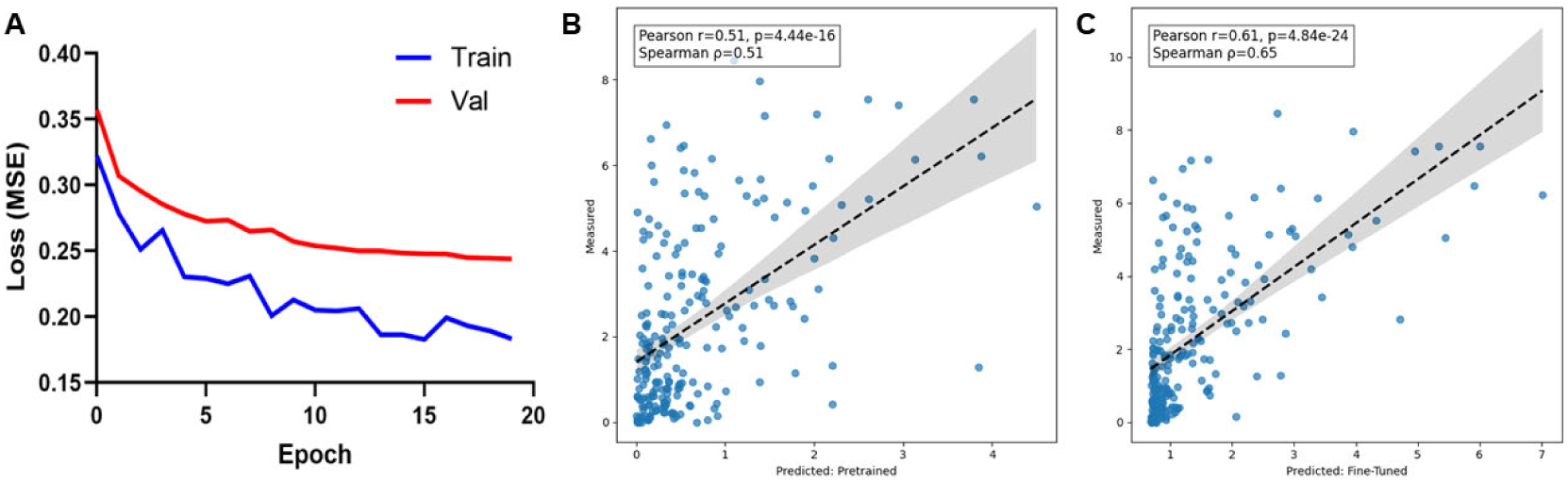
Fine-tuning improves gene expression prediction in macrophages. (A) Training and test loss curves over 20 epochs of model fine-tuning. (B–C) Predicted versus measured RNA expression values for 230 transcripts induced by TNF*𝛼* in macrophages, shown for the pretrained model (B) and the fine-tuned model (C).

To interpret the model’s predictions and identify candidate regulatory elements, we applied a widely adopted backpropagation-based technique, DeepLIFT (Shrikumar et al., 2019), to compute nucleotide-resolution importance scores, capturing the contribution of each base to splicing of the retained intron. Attribution scores were computed for each intron along with sequence within the model input window (524 kb). In these scores, larger positive values indicate nucleotides that the model predicts as contributing to intron retention (i.e., delaying splicing), whereas negative values indicate nucleotides pre-dicted to be associated with splice completion. We analyzed these scores using modiscolite, which clusters recurring high-importance patterns into position weight matrices (PWMs) representing putative regulatory motifs (Shrikumar et al., 2020). To assess motif enrichment among delayed introns, we focused on the 50 slowest-splicing introns as ranked by area under the CoSI-time curve (Fig. S7C). Motif occurrences were identified using a conventional motif scanning approach, Finding Individual Motif Occurrences (FIMO) (Grant et al., 2011), and enrichment was calculated relative to either the remaining introns in our dataset or all introns in the transcriptome.

We compared the frequency of putative regulatory motifs in the 50 slowest-splicing introns to their frequency across all introns in the murine transcriptome, highlighting both the most enriched motifs overall and specific examples of their genomic locations within delayed introns. Among these, a GA-repeat–like motif showed enrichment in the slow-splicing set (FIMO, 80.0% vs 55.2%) (Fig. 6A). Representative instances of this motif were also recovered from the original source seqlets used to generate the PWM by modiscolite, providing further support for its regulatory relevance (Fig. 6B–D). These findings illustrate how sequence-to-expression modeling, combined with attribution-based motif discovery, can uncover non-canonical cis-regulatory elements that may influence splicing efficiency during inflammatory gene activation.

**Figure 6.**
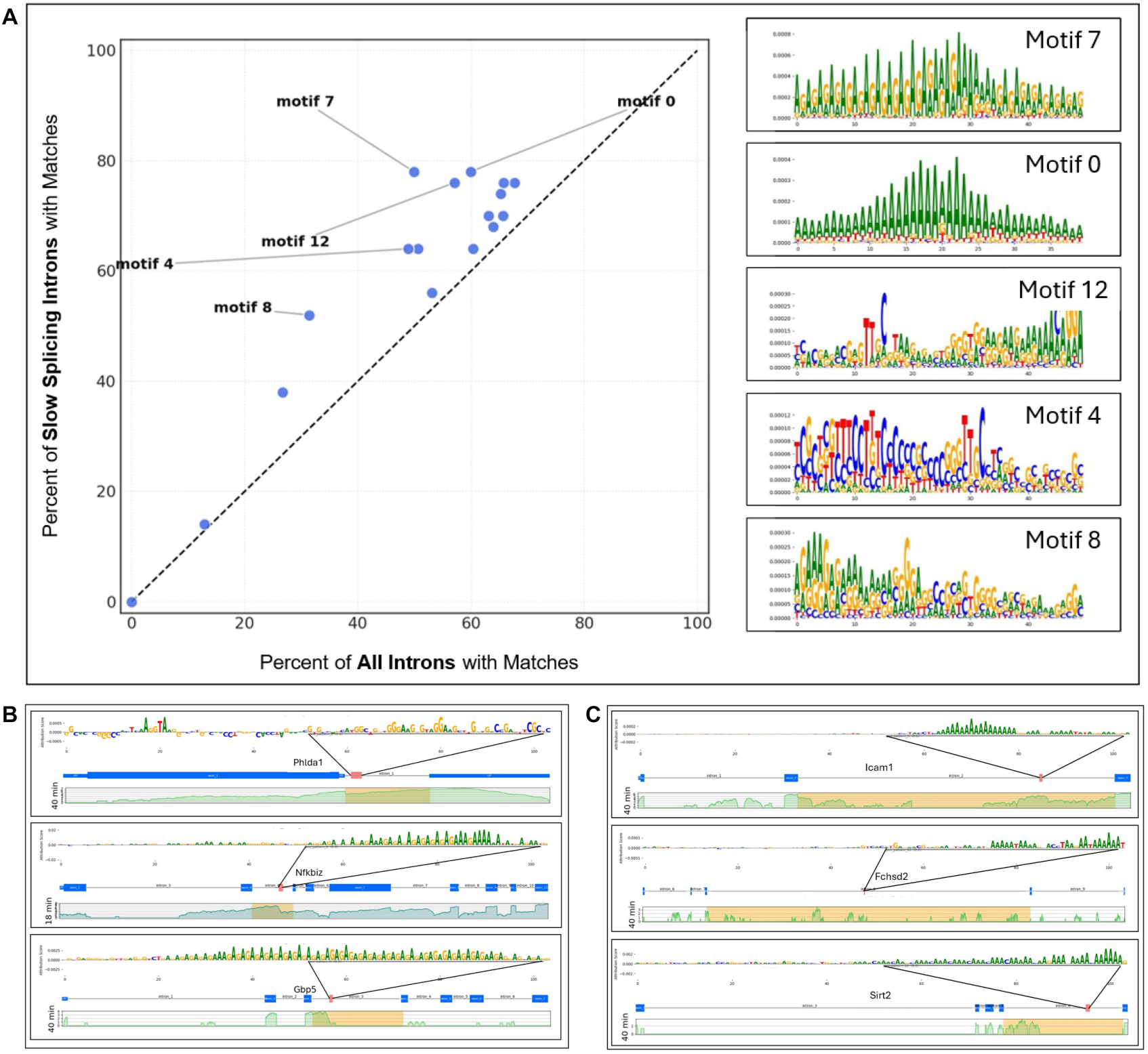
Putative regulatory sequences are enriched in slow-splicing introns. (A) Scatter-plot showing percent representation of position weight matrices (PWMs) scanned across slow-splicing introns versus all introns genome-wide using FIMO. Sequence logos of the top five enriched PWMs are shown to the right. (B–D) Attribution plots highlighting GA-rich sequences (source seqlets for enriched PWMs) and their locations mapped to gene schematics for GA-rich motif (B) and A-rich motif (C). Corresponding RNA-seq tracks are shown for each gene.

## Discussion

In this study we sought to understand splicing kinetics of the large number of genes that comprise the inflammatory response and to assess whether splicing itself might play a regulatory role in inflammation. We developed a targeted sequencing strategy, purifying transcripts containing each gene’s terminal exon. This approach allowed us to sequence the 1,024 introns within inflammatory genes and permitted direct assessment of the structures of nearly-completed transcripts. We found considerable heterogeneity in splicing efficiency among these introns. In studying evolutionarily conserved weak 5^′^ splice donors, we have isolated one cause of slow appearance of mRNA following a pulse of stimulus; many other slowly spliced introns without such sequences were also identified in this study and suggest other regulatory mechanisms may be responsible.

Crucially, the hybrid capture approach averts a common ambiguity in analyzing splicing kinetics of not being able to differentiate completed pre-mRNA from nascent transcripts during an induction pulse—this often leads to an overestimation of the unspliced status of early introns and complicates quantification of splicing kinetics. To the contrary, we rarely found genes containing unspliced first introns. This was true of chromatin-associated RNA and of whole-cell RNA after inhibition of transcription with actD. These effects are consistent with the emerging model of co-transcriptional pre-mRNA splicing, where the splicing machinery has been suggested to lag 3–5 kb behind the polymerase. Indeed, several recent global studies of RNA splicing bolster the claim that much pre-mRNA is spliced co-transcriptionally: 74% (yeast) or 75–84% (human) of introns are found to be at least 50% spliced by the time of transcription termination in several other studies (Tilgner et al., 2012; Carrillo Oesterreich et al., 2010; Ameur et al., 2011; Brugiolo et al., 2013; Girard et al., 2012; Khodor et al., 2011). Surprisingly, this ∼80% figure remains constant whether total RNA or chromatin-associated RNA is measured, implying that our choice to analyze chromatin-associated RNA does not significantly overrepresent splicing intermediates.

We found that most introns are spliced very efficiently, appearing and disappearing as a rapid dip of CoSI immediately following induction, returning to a CoSI of ∼0.95 within minutes after induction. Notably, the distribution of CoSI values of the entire dataset (Fig. 2) was very broad. Though most introns spliced immediately, there were several “bottleneck introns.” In order to determine more specifically the rates of slowly splicing introns, studies employing actD to stall transcription and examine intron splicing half-lives corroborated the idea that there is tremendous intron-to-intron heterogeneity. Most delayed introns ultimately reached higher CoSI values over the time course, consistent with completion of splicing rather than stable, long-term retention (see Figs. 2–3). Whereas most introns spliced within 20–40 s, some were delayed significantly (upwards of 5 minutes). Of note, however, is that our 20–40 s rate of splicing is somewhat at odds with other figures in the literature of 8–10 minutes for intron excision (Pandya-Jones et al., 2013) after a washout of the drug D-ribofuranosylbenzimidazole. There is some debate as to the perturbative role of actD in splicing, with one report observing that splicing intermediates in the context of the MS2 reporter system are prematurely liberated from chromatin upon actD treatment. Even in this case, the rapid actD-based rates are likely underestimating even faster kinetics if one considers that the co-transcriptional splicing machinery targets chromatin mRNA faster than released mRNA (Martin et al., 2013). However, even in the absence of actD, stimulation revealed that most of *I𝜅B𝛼*’s introns are spliced in less than two minutes when one takes into account the “dip” in CoSI due to induction and the time to reach steady CoSI levels (Fig. 2). While the terminal intron of *I𝜅B𝛼* appears to have a longer half-life, this unique feature of terminal introns is consistent with prior studies (Carrillo Oesterreich et al., 2010).

In testing gene expression differences in bottleneck introns among introns with poor splice sites that are also evolutionarily conserved, we found that steady-state levels of reporter proteins were upregulated when the 5^′^ splice donor sequence was mutated to the consensus sequence ‘GTAAG’ in all cases but one. Attenuated U1 binding provides a mechanistic insight for bottleneck introns that were chosen for their weak 5^′^ splice donors. This implies that at the level of splicing, either due to delays in expression or perhaps degradation due to delayed expression, significant differences in gene expression arise from small differences in nucleotide sequence. These reporter assays measure steady-state expression influenced by intron sequence; while consistent with changes in splicing efficiency, other post-transcriptional mechanisms may be at work to preclude these introns from efficient splicing. We also find many slowly spliced introns not explained by weak 5^′^ splice donors. In some cases, we find multiple bottleneck introns per gene, as is the case of *IRF7*, where one bottleneck (intron 5) was attributable to an evolutionarily conserved weak 5^′^ splice donor while another (intron 7) was observed experimentally but of unknown cause. This may be due to any of a number of potential mechanisms that may also serve in tuning the speed of splicing: cis-regulatory protein recruitment, 3^′^ splice acceptor sequence or other sequence elements, or alterations of RNA polymerase speed or chromatin marks or three-dimensional gene structure.

Central to our inquiry is the enigmatic nature of these bottlenecks remaining in physiologically critical genes, often evolutionarily conserved, and yet intrinsically mediating an inefficiency in gene expression. Importantly, the conservation of these weak donor sites suggests they confer a regulatory advantage—such as fine-tuning of expression timing or transcript abundance—rather than representing neutral or deleterious features merely tolerated by selection. We posit that the gene expression changes that are shown in bone marrow-derived macrophages offer a regulatory strategy to slow up and maybe restrict expression of genes in a manner dependent on the composition of mRNA processing factors in the cell (“the splicing landscape”), the cell type, or the stimulus type in question. Recent studies have demonstrated global changes in intron retention preferences in B cell lymphomagenesis and granulocyte differentiation (Koh et al., 2015). In a similar manner, we suggest that selection of splicing and kinetics of splicing might allow a previously unappreciated level of specificity to gene expression decisions in cells presented with an inflammatory stimulus (57–63).

Induction with TNF is a particularly favorable situation because many of the genes we examined were up-regulated in their transcription within 4–6 minutes of adding inducer (Fig. S2B), allowing examination of large numbers of pre-mRNA transcripts. This, in concert with the hybrid capture approach that provides a large number of junctional sequencing reads, has permitted unique insight into the kinetics of splicing of mature transcripts and revealed surprising heterogeneity. We suggest that this methodology and analysis could have wider applicability for other gene induction situations.

Despite the promise of deep learning models, challenges remain in translating their predictions into biological insight. While performance on benchmark tasks continues to improve—often surpassing traditional motif-finding tools—generalization across conditions and cell types is still constrained by the scope of training data. Interpretability is another major hurdle, particularly for genomic sequences. Interpretable machine learning methods can highlight sequence elements that influence model predictions, but connecting these regions to underlying biological mechanisms often demands extensive experimental validation. In the case of RNA splicing, complexity of overlapping regulatory layers—such as RNA secondary structure, co-transcriptional dynamics, and nuclear export—can make it difficult to disentangle causal sequence features from correlated signals. Nonetheless, sequence-to-expression modeling provides a powerful framework for identifying sequence features correlated with delayed splicing and for guiding experimental discovery of novel regulators through fine-tuning and perturbation-informed learning.

Our deep learning–based analysis further supports the notion that splicing efficiency is shaped by a broad array of cis-regulatory features, many of which fall outside canonical splice site motifs. By leveraging a sequence-to-expression transformer trained on multimodal regulatory data, we identified non-canonical sequence motifs enriched in the slowest-splicing introns—motifs that may act as silencers or delay elements involved in splicing. These results offer a complementary, agnostic perspective to our mutational analyses, and suggest that intronic bottlenecks can arise from diverse sequence architectures. The enrichment of these motifs in bottleneck introns—independent of donor site strength—points to additional layers of splicing control that may be particularly relevant in rapid-response transcriptional programs like inflammation. We propose that such regulatory elements may encode a form of temporal tuning, modulating transcript availability through fine control of intron excision. More broadly, our work demonstrates that interpretable machine learning can uncover latent regulatory features that elude traditional sequence analysis, advancing efforts to decode the logic of splicing regulation in a cell-type or stimulus-specific context.

## Methods

### Cells

C56BL6/J mice were sacrificed via CO_2_ euthanasia and sterilized with 70% ethanol. Femur and tibia bones were harvested and stripped of muscle tissue. Bone marrow cells were resuspended in 20 mL of fresh DMEM. 2.5 ×10^6^ bone-marrow cells were plated in a 15-cm dish in 20 mL of BMDM media (DMEM, 20% FBS, 30% L929 conditioned media, and 1% Pen/Strep) and grown at 5% CO_2_ and 37^◦^C. BMDM media was completely replaced on day 3 as well as a supplemental addition of 5 mL L929 conditioned media on day 5.

### RNA fractionation

RNA was fractionated into cytoplasmic, nucleoplasmic, and chromatin-associated pools as previously described (Bhatt et al., 2012) with modifications. Confluent 15 cm dishes of mature BMDMs were scraped into 400 *𝜇*L cold NP-40 lysis buffer (10 mM Tris-HCl pH 7.5, 0.08% NP-40, 150 mM NaCl) and layered onto a 1 mL sucrose cushion (10 mM Tris-HCl pH 7.5, 150 mM NaCl, 24% w/v sucrose). Samples were centrifuged at 13,000 rpm for 10 min at 4^◦^C. The supernatant (cytoplasmic fraction) was mixed with 3 volumes of 100% ethanol and 2 volumes of buffer RLT (4 M GuSCN, 0.1 M *𝛽*-mercaptoethanol, 0.5% N-lauroylsarcosine, 25 mM Na-citrate, pH 7.2) and stored at –80^◦^C.

The nuclear pellet was resuspended in 200 *𝜇*L cold glycerol buffer (20 mM Tris-HCl pH 7.5, 75 mM NaCl, 0.5 mM EDTA pH 8.0, 50% glycerol, 0.85 mM DTT) and lysed with an equal volume of nuclear lysis buffer (20 mM HEPES pH 7.5, 7.5 mM MgCl_2_, 0.2 mM EDTA pH 8.0, 1 M urea, 1% NP-40). After vortexing and centrifugation (14,000 rpm, 5 min, 4^◦^C), the supernatant (nucleoplasmic fraction) was processed as above.

The remaining chromatin pellet was hydrated in 1× PBS and dissolved in 500 *𝜇*L TRIzol reagent at 50^◦^C with intermittent vortexing. After phase separation with 100 *𝜇*L chloroform, the aqueous phase was recovered and processed as above. All RNA fractions were purified using the Qiagen RNeasy protocol and eluted in nuclease-free water. Typical yields were 300–500 ng/*𝜇*L for cytoplasmic RNA, 100–250 ng/*𝜇*L for nucleoplasmic RNA, and 300–500 ng/*𝜇*L for chromatin-associated RNA. RNA was DNase-treated (TURBO DNase, Thermo Fisher) and stored at –80^◦^C.

### Template-switch reverse transcription

One microgram of RNA was mixed with 1 *𝜇*M oligo(dT)_30_ (5^′^-AAGCAGTGGTATCAACGCAGAGTACT_30_-3^′^), heated to 80^◦^C for 2.5 min, and snap-cooled on ice. A 10 *𝜇*L reversetranscription mix containing 10 *𝜇*M template-switch oligo (5^′^-AAGCAGTGGTATCAACGCAG AGTACACArGrGrG-3^′^), 20 mM DTT, 2× First-Strand Buffer (Invitrogen), 1 mM dNTPs, 40 U Murine RNase Inhibitor (NEB), and 200 U SuperScript II (Invitrogen) was added. Reactions were incubated sequentially at 42^◦^C (30 min), 45^◦^C (30 min), and 50^◦^C (10 min), then heat-inactivated at 80^◦^C (10 min).

RNA templates were degraded by adding NaOH (final 0.1 M) and EDTA (5 mM) and heating to 70^◦^C for 10 min, followed by neutralization with HCl. cDNA was purified using 2× Sera-Mag carboxylate-modified magnetic beads (GE Healthcare) with standard PEG/ethanol washes and eluted in nuclease-free water.

### Hybrid capture probe design and synthesis

Biotinylated RNA probes were designed against the terminal exons of inflammatory genes (see Supplemental Table 2). For each gene, 100-bp overlapping oligonucleotides were synthesized (CustomArray Inc.) and pooled into nine expression-matched subgroups. Each subpool was PCR-amplified to append a T7 RNA polymerase promoter and transcribed in vitro using the AmpliScribe T7 Biotin Flash Kit (Epicentre). Purified RNA probe subpools were combined in weighted ratios (A–I) to normalize capture representation across expression levels as below:

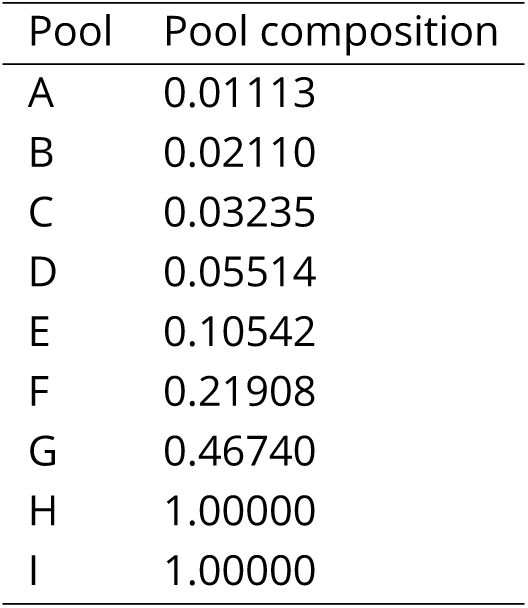

| Pool | Pool composition |
| --- | --- |
| A | 0.01113 |
| B | 0.02110 |
| C | 0.03235 |
| D | 0.05514 |
| E | 0.10542 |
| F | 0.21908 |
| G | 0.46740 |
| H | 1.00000 |
| I | 1.00000 |

### cDNA hybrid capture and elution

Biotinylated probes were hybridized to cDNA at 74^◦^C for 4.5 min, followed by addition of 2× hybridization buffer (1 M LiCl, 40 mM Tris-HCl pH 7.5, 20 mM EDTA pH 8.0, 4 M urea, 0.5% Triton X-100, 1% SDS, 0.2% Na-deoxycholate). Hybridization continued 30 min at 70^◦^C. Streptavidin BioMag beads (0.3 mg) were washed and incubated 20 min at 70^◦^C to capture cDNA–probe complexes. Beads were washed twice with 1× HYB, once each with Wash 4 and Wash 5 buffers, and eluted in 35 *𝜇*L base elution buffer (125 mM NaOH, 10 mM EDTA pH 8.0, 10 mM Tris-HCl pH 7.5) at 74^◦^C for 5 min. The eluate was neutralized and purified with 1× Sera-Mag beads and eluted in 45 *𝜇*L and stored at –80^◦^C.

### Determining the efficiency of cDNA pulldown

Pulldown efficiency was evaluated by qPCR to quantify enrichment of target transcripts and depletion of background RNA. qPCR reactions (KAPA SYBR 2× Master Mix) compared preand post-pulldown cDNA at a 2:1 ratio. Primers:

- L32 (background control): F 5^′^-AAGCGAAACTGGCGGAAAC-3^′^; R 5^′^-TAACCGATGTTG GGCATCAG-3^′^
- NF-*𝜅*BIA (spliced exon 5–6 junction): F 5^′^-ACGGAGTCAGAATTCACAGAGG-3^′^; R 5^′^-CA CAAAGACAACAGCCGAATC-3′

Cycle-threshold (Ct) values were used to calculate ΔCt between L32 and NF-*𝜅*BIA. Successful pulldowns typically showed ΔL32 = –7 to –9 cycles and ΔNF-*𝜅*BIA = –2 to –4, corresponding to ΔL32/ΔNF-*𝜅*BIA *>* 2.0.

### Post-pulldown cDNA amplification

Pulldown cDNA was amplified prior to tagmentation. PCR reactions contained Q5 High-Fidelity 2× Master Mix (NEB), 1 *𝜇*M primer (5^′^-AAGCAGTGGTATCAACGCAGAGTACT-3^′^), and ∼5% of the pulldown reaction. Cycling: 95^◦^C (2 min) → 20–25 cycles of 95^◦^C (30 s), 62.5^◦^C (30 s), 72^◦^C (150 s) → 72^◦^C (5 min). PCR products were purified (0.9× Sera-Mag) and eluted in 25 *𝜇*L H_2_O. Concentrations were determined using a Qubit HS dsDNA Assay.

### Tagmentation of cDNA libraries with Tn5 transposase

Tn5 transposase was purified as in Picelli et al., *Genome Res.* (2014) and pre-loaded with hybridized adapter oligos (Tn5MErev 5^′^-[phos]CTGTCTCTTATACACATCT-3^′^; Tn5ME-A 5^′^-T CGTCGGCAGCGTCAGATGTGTATAAGAGACAG-3^′^; Tn5ME-B 5^′^-GTCTCGTGGGCTCGGAGATGTGTATAAGAGACAG-3^′^). Tagmentation reactions contained ∼40 ng amplified cDNA, 0.2 *𝜇*L Tn5, 5% PEG8000, 10 mM TAPS (pH 8.5), 5 mM MgCl_2_, incubated 10 min at 55^◦^C. SDS (0.02%) was added and incubation continued 10 min to inactivate Tn5. Reactions were 1.4× Sera-Mag purified and eluted in 20 *𝜇*L H_2_O.

### Library barcoding and sequencing

Tagmented libraries were PCR-amplified with paired barcode oligos using Q5 High-Fidelity 2× Master Mix (NEB). Cycling: 72^◦^C (3 min), 98^◦^C (2 min), then 25 cycles of 98^◦^C (10 s), 63^◦^C (30 s), 72^◦^C (30 s), and 72^◦^C (5 min). Libraries were double-purified (1.0×/1.4× Sera-Mag), quantified by Qubit HS dsDNA assay, pooled equimolarly, repurified, and analyzed with an Agilent Bioanalyzer 2100. Sequencing was performed on an Illumina HiSeq 2500 (50 bp, single-end mode).

### RNA-seq alignment and analysis

Single-end 50 bp reads were aligned to the mm10 genome using STAR. Junctions were retained only if both exonic and intronic segments were ≥3 bp. Using pysam, splice and intron junctions were classified as types a, b, or c, and Completion of Splicing Index (CoSI) values were computed using custom Python scripts. All analyses were performed in Python and R. Sequencing data are being deposited to NCBI GEO, and all analysis code will be made available on GitHub.

### Deep Learning Analysis of Intron Regulatory Features

#### Model architecture and initialization

We used Borzoi, a transformer-based sequence-to-function model trained to predict steady-state RNA abundance from genomic DNA sequence. The pretrained model captures general patterns of cis-regulatory architecture by integrating multi-omic training targets (CAGE, ATAC-seq, and ChIP-seq tracks). However, because Borzoi is optimized for baseline expression rather than stimulus-responsive contexts, we adapted it to our macrophage TNF*𝛼* time course through targeted fine-tuning.

#### Fine-tuning on TNF*𝛼*-stimulated macrophage data

We generated genome-wide RNA-seq coverage tracks from bone marrow–derived macrophages (BMDMs) collected at 18, 20, and 30 minutes post-TNF*𝛼* induction—timepoints representative of dynamic intron retention. These coverage values served as supervised training targets to refine the model’s prediction of RNA abundance under inflammatory conditions. Fine-tuning was performed for 20 epochs using the Adam optimizer (learning rate = 1×10^−6^, mean-squared-error loss). To evaluate generalization, chromosomes 10 and 11 were withheld for validation and testing, respectively. Model loss steadily decreased over training for both sets, and the correlation between predicted and observed expression improved from Pearson *𝑟* = 0.51 (*𝑝* = 4.44×10^−16^) for the pretrained model to *𝑟* = 0.61 (*𝑝* = 4.84×10^−24^) after fine-tuning (Fig. 5A–B).

#### Feature attribution and motif discovery

To interpret the fine-tuned model’s predictions, we applied DeepLIFT to compute nucleotide– resolution importance scores, quantifying each base’s contribution to the predicted RNA signal within a 524 kb genomic window centered on each intron of interest. Positive importance scores indicate sequence positions contributing to higher predicted intronic RNA signal (i.e., delayed splicing), whereas negative scores correspond to features predictive of splice completion.

Importance profiles were analyzed using modisco-lite, which clusters recurrent high-importance patterns across examples into position weight matrices (PWMs) representing putative regulatory motifs. Enrichment of discovered motifs was assessed using FIMO, scanning the 50 (or 150) slowest-splicing introns—ranked by area under the CoSI–time curve—against all annotated murine introns (GENCODE M23). For targeted motif discovery, attributions input to modisco-lite were masked under different configurations to restrict pattern identification to either intronic or exonic regions adjacent to slow-splicing introns.

#### Motif enrichment and visualization

Relative motif frequencies were plotted as a scatter of percent representation in slow- versus all-intron backgrounds. Sequence logos of the top five enriched motifs were visualized alongside genomic examples where GA-rich and A-rich motifs coincided with regions of high attribution in DeepLIFT profiles (Fig. 6A–D).

## Acknowledgements

The authors would like to thank Alex Shishkin and Mitchell Guttman (Dept. of Biology, Caltech) for assistance with hybrid capture strategy design; and Ann-Jay Tong, Stephen Smale, Doug Black, and Amy-Pandya Jones (Dept. of Biology, University of California, Los Angeles) for insights and advice; and Sergei Manakov, Evelyn Stuwe, Dubravka Pezic, Igor Antoshechkin, Sagar Damle, and Alok Joglekar (Dept. of Biology, California Institute of Technology) for experimental and computational assistance. This work was funded from a grant from NIH and from an endowment provided by the Raymond and Beverly Sackler Foundation. Research reported in this publication was supported by the National Institute of General Medical Sciences (NIGMS) of the National Institutes of Health under award number P20GM125498.

## Author Contributions

J.S. Dearborn, Conceptualization, Investigation, Data curation, Software, Formal analysis, Visualization, Methodology, Writing – review & editing. L. Frankiw, Conceptualization, Investigation, Resources, Methodology, Validation, Writing – review & editing. D.W. Limoge, Data curation, Formal analysis, Visualization, Writing – review & editing. C.H. Burns, L. Vlach, P. Turpin, T. Kirch, Z.D. Miller, W. Dowell, S. Languon, Y. Garcia-Flores, Investigation, Data curation, Validation, Writing – review & editing. R.C. Cockrell, Resources, Supervision, Project administration, Writing – review & editing. D. Baltimore, Conceptualization, Supervision, Funding acquisition, Project administration, Writing – review & editing. D. Majumdar, Conceptualization, Supervision, Funding acquisition, Project administration, Methodology, Writing – review & editing.

## Supplementary Figure Legends

**Supplementary Figure 1.**
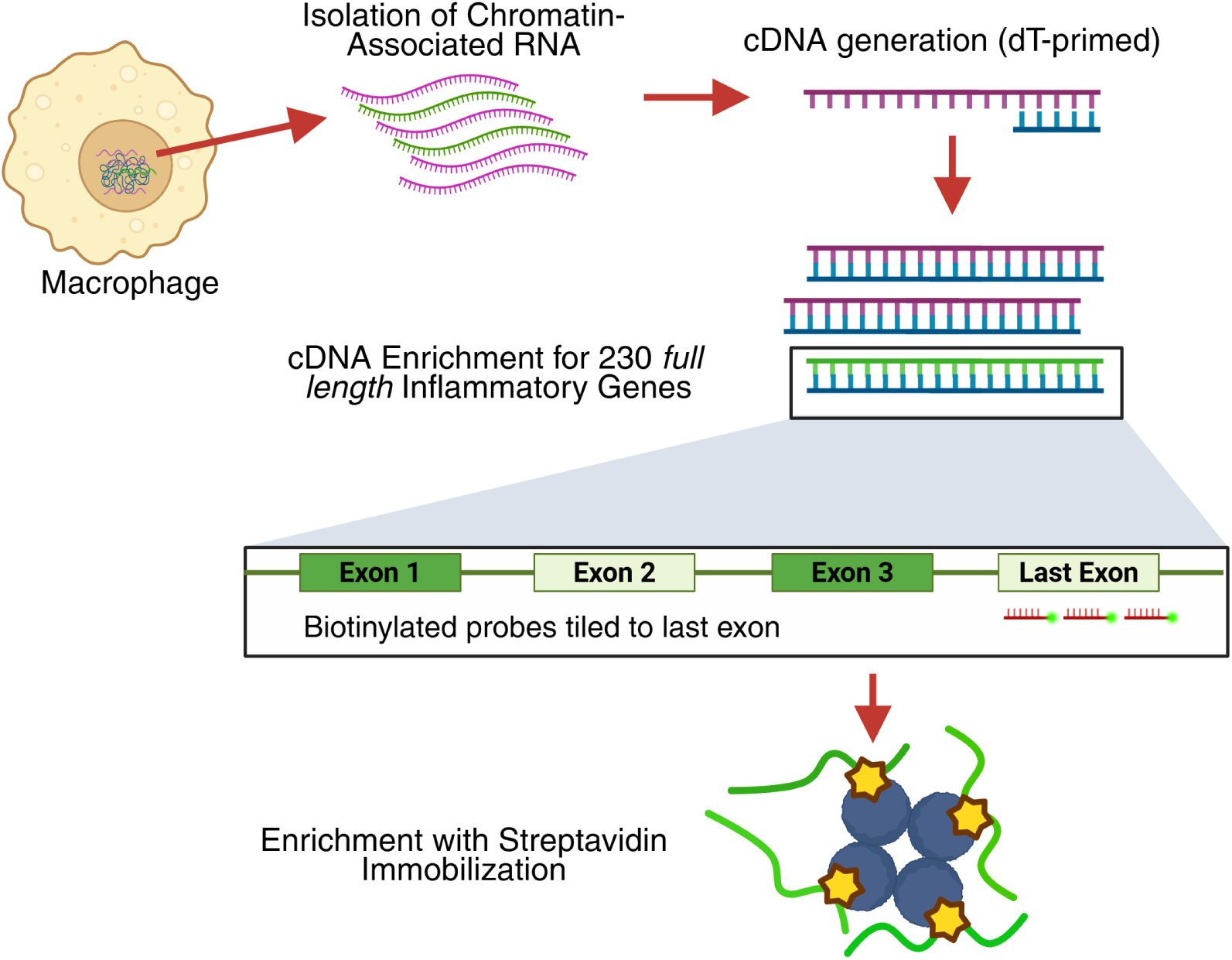
Hybrid capture strategy for isolating chromatin-associated inflammatory transcripts. RNA was purified from chromatin-associated bone marrow– derived macrophages (BMDMs), and cDNA was generated using oligo(dT) priming to enrich for polyadenylated transcripts. Biotinylated RNA oligonucleotides complementary to the terminal exons of inflammatory genes were hybridized to the cDNAs, allowing for selective enrichment of these transcripts via streptavidin bead capture.

**Supplementary Figure 2.**
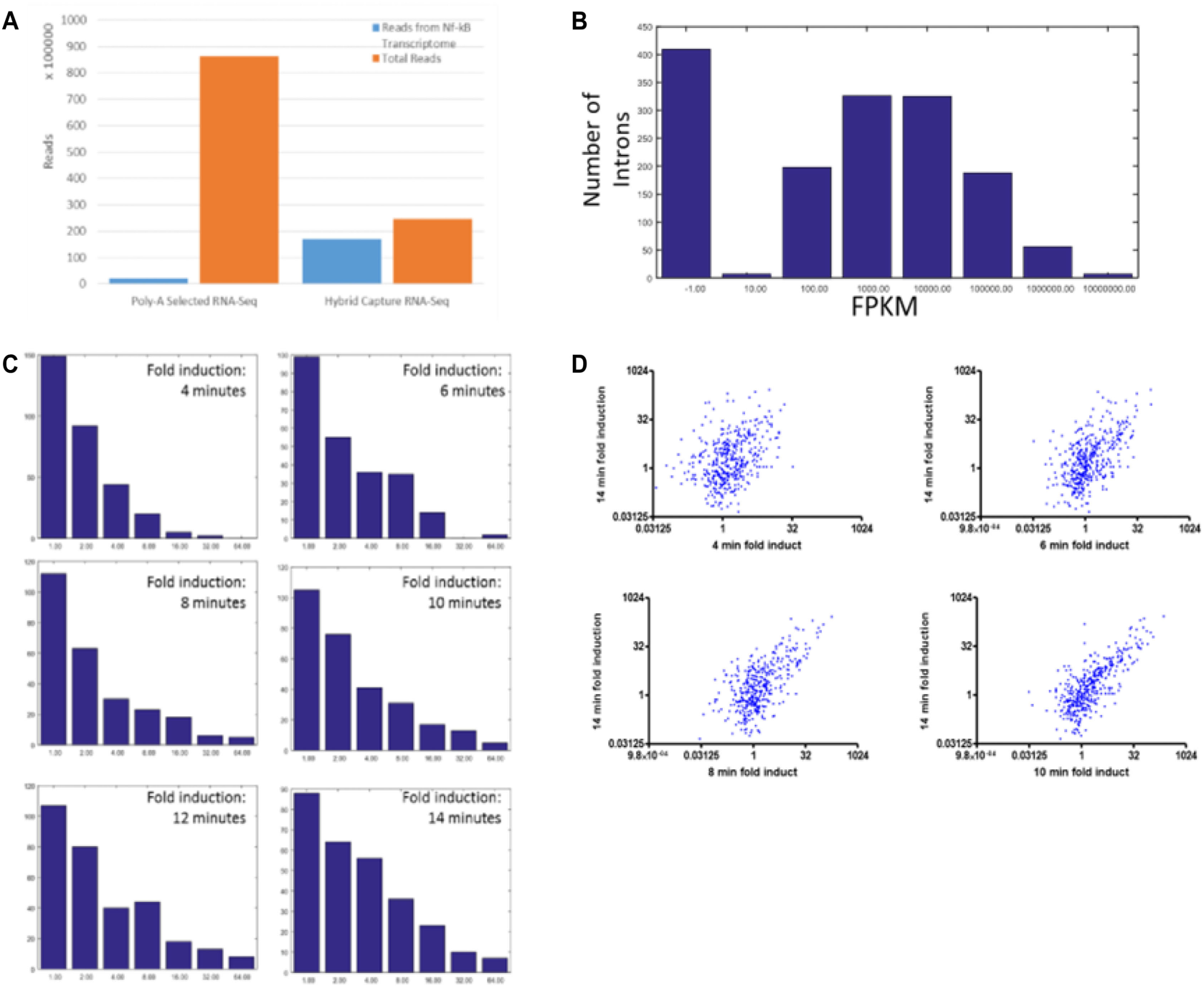
Hybrid capture enriches NF-*𝜅*B–responsive transcripts and yields robust intron coverage. (A) Fraction of sequencing reads corresponding to NF-*𝜅*B– responsive genes (blue) in hybrid-captured versus poly(A)-selected RNA (orange), showing substantial enrichment after hybrid capture. (B) Histogram of read counts per intron at the 6-min TNF induction time point. Reads were detected for 1,024 introns out of 1,508 targeted introns, with undetected introns largely corresponding to transcripts induced at later time points (*>*60 min). (C) Distribution of intron fold induction across time points reveals that many NF-*𝜅*B target genes begin to show induction as early as 4 min post-TNF stimulation, with both the number of induced introns and magnitude of induction increasing markedly by 14 min. (D) Scatterplot comparing induction at 4 min versus 14 min for individual introns demonstrates that most early-induced genes are further upregulated at later time points.

**Supplementary Figure 3.**
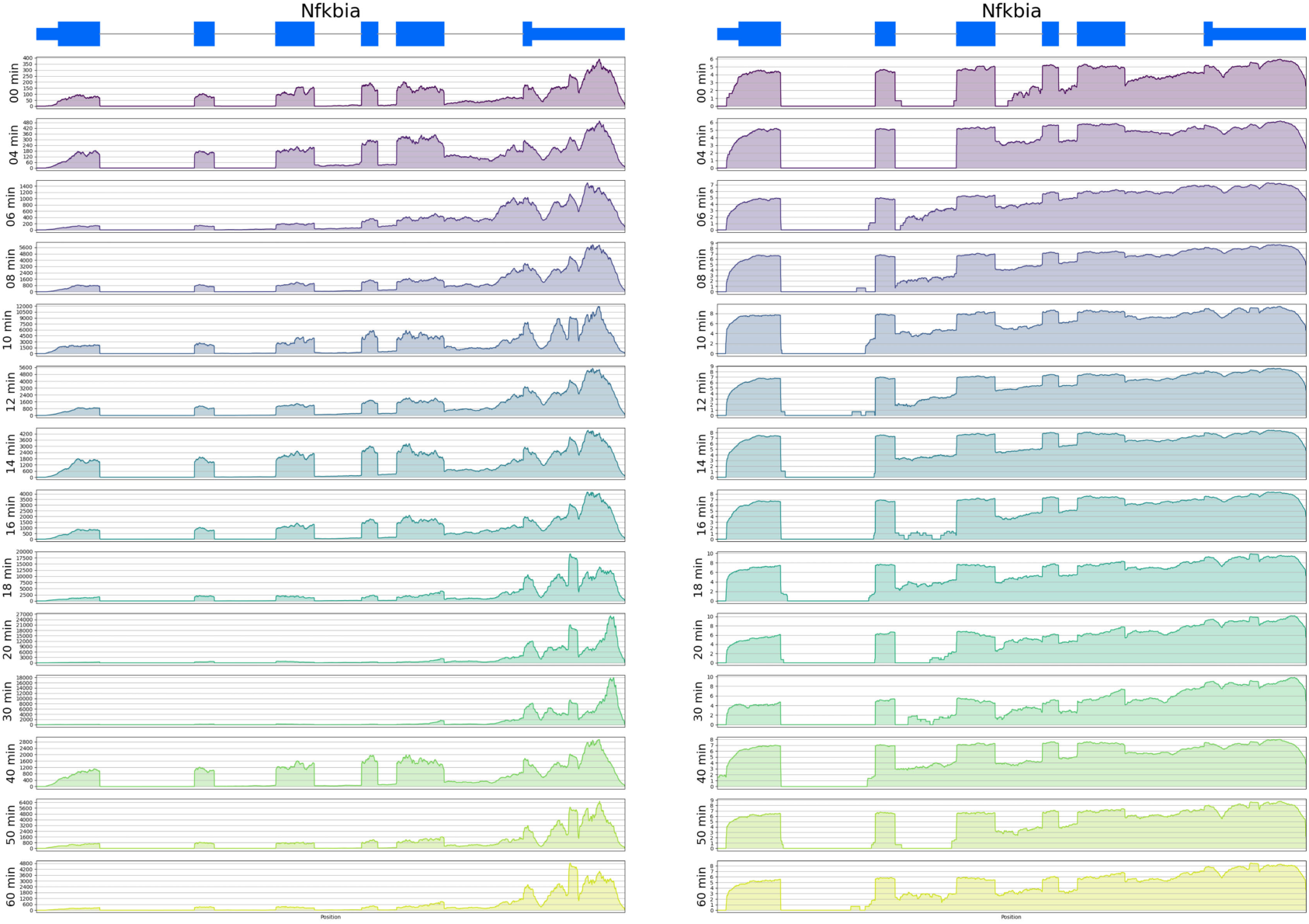
Gene track visualization of *Nfkbia* induction dynamics. Genome browser tracks show *Nfkbia* (IKB*𝛼*) transcript induction over time in chromatin-associated, hybrid-captured RNA. Signal intensity is displayed on a linear scale (left) normalized to the maximum height at 20 min, and on a log scale (right) normalized to the maximum at each time point.

**Supplementary Figure 4.**
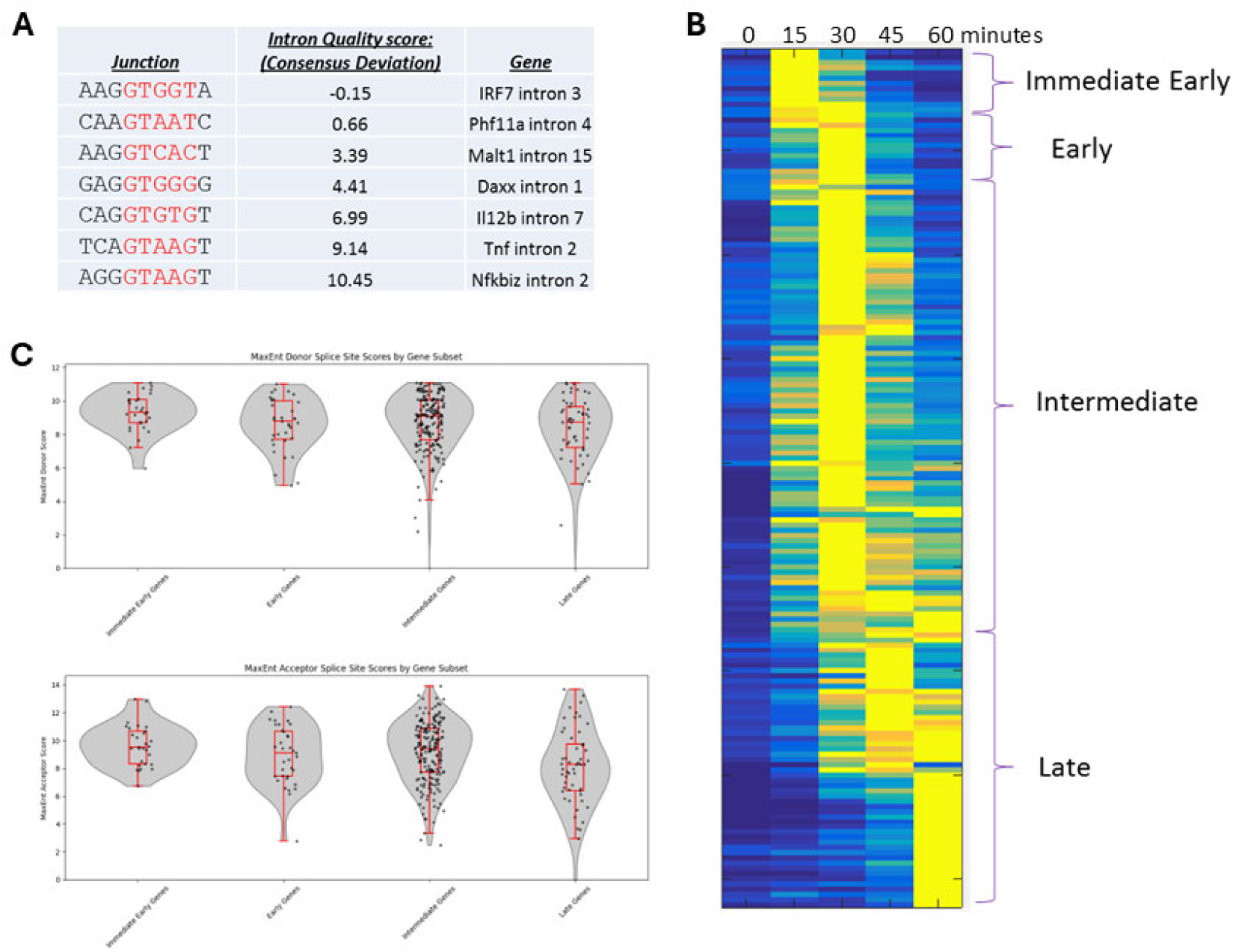
Splice site strength and expression kinetics among NF-*𝜅*B–induced genes. (A) Representative introns from selected genes were scored for 5^′^ splice donor strength based on similarity to the canonical “GTAAG” motif; introns with weaker matches (e.g., *Irf7* intron 3) received lower scores. (B) NF-*𝜅*B–induced genes display well-characterized variability in expression kinetics (RNA-seq data from Reference 25). Heatmap shows temporal expression profiles of NF-*𝜅*B target genes following lipid A stimulation, categorized into immediate-early, early, and later expression groups. (C) All introns in the NF-*𝜅*B transcriptome were scored using MaxEntScan for both 5^′^ splice donor (top) and 3^′^ splice acceptor (bottom) sequences, stratified by their gene expression group.

**Supplementary Figure 5.**
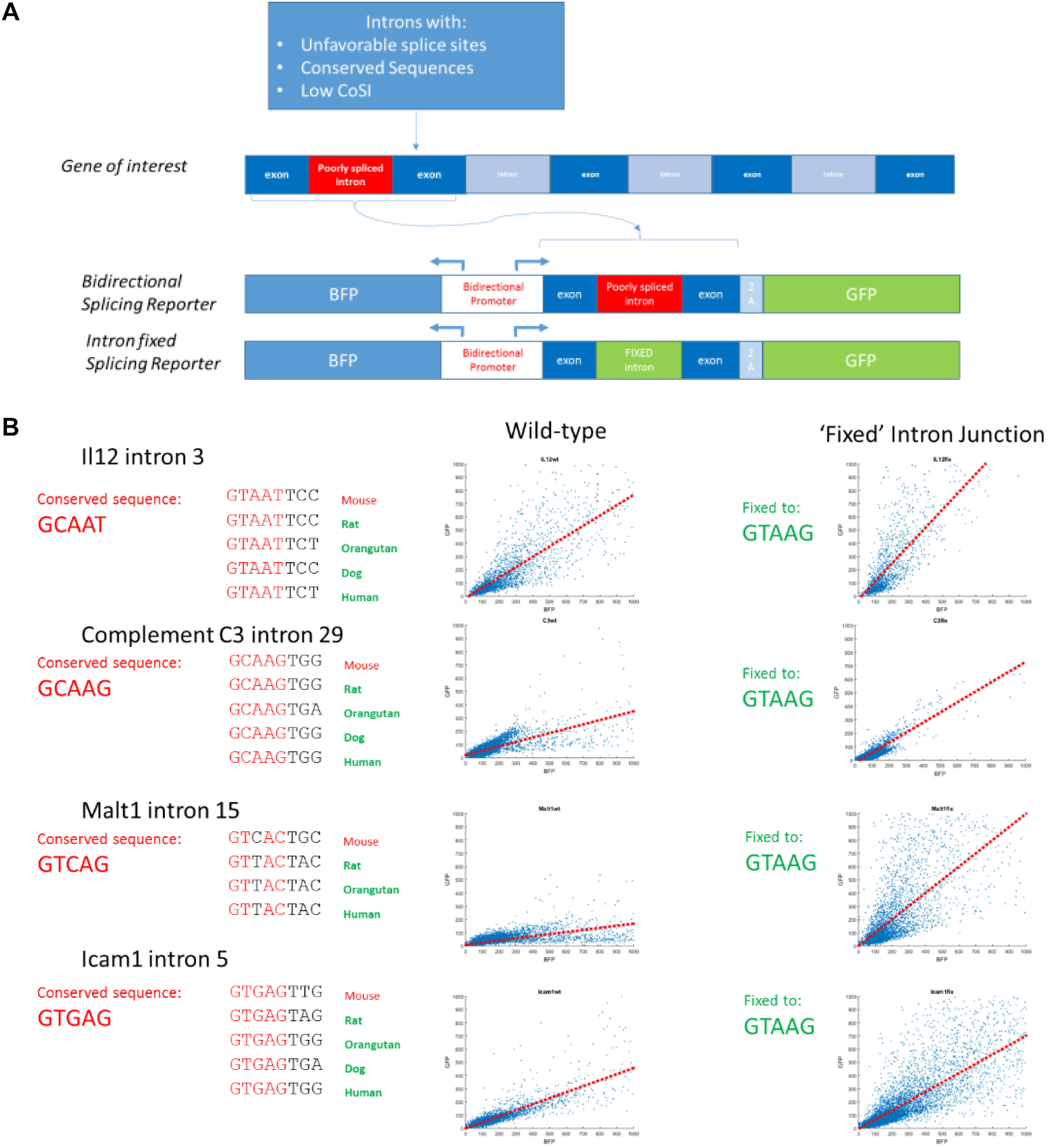
Reporter assay design for assessing intron splicing efficiency. (A) Schematic of the bidirectional reporter assay. Individual introns were cloned into a bidirectional promoter context together with their flanking exons, positioned upstream of a self-cleaving 2A peptide and GFP reporter. In the opposite transcriptional direction, a BFP reporter served as a transcriptional control. (B) Evolutionarily conserved weak 5^′^ splice donors were “repaired” to the canonical GTAAG sequence within this reporter construct, and splicing efficiency was quantified by flow cytometry (FACS).

**Supplementary Figure 6.**
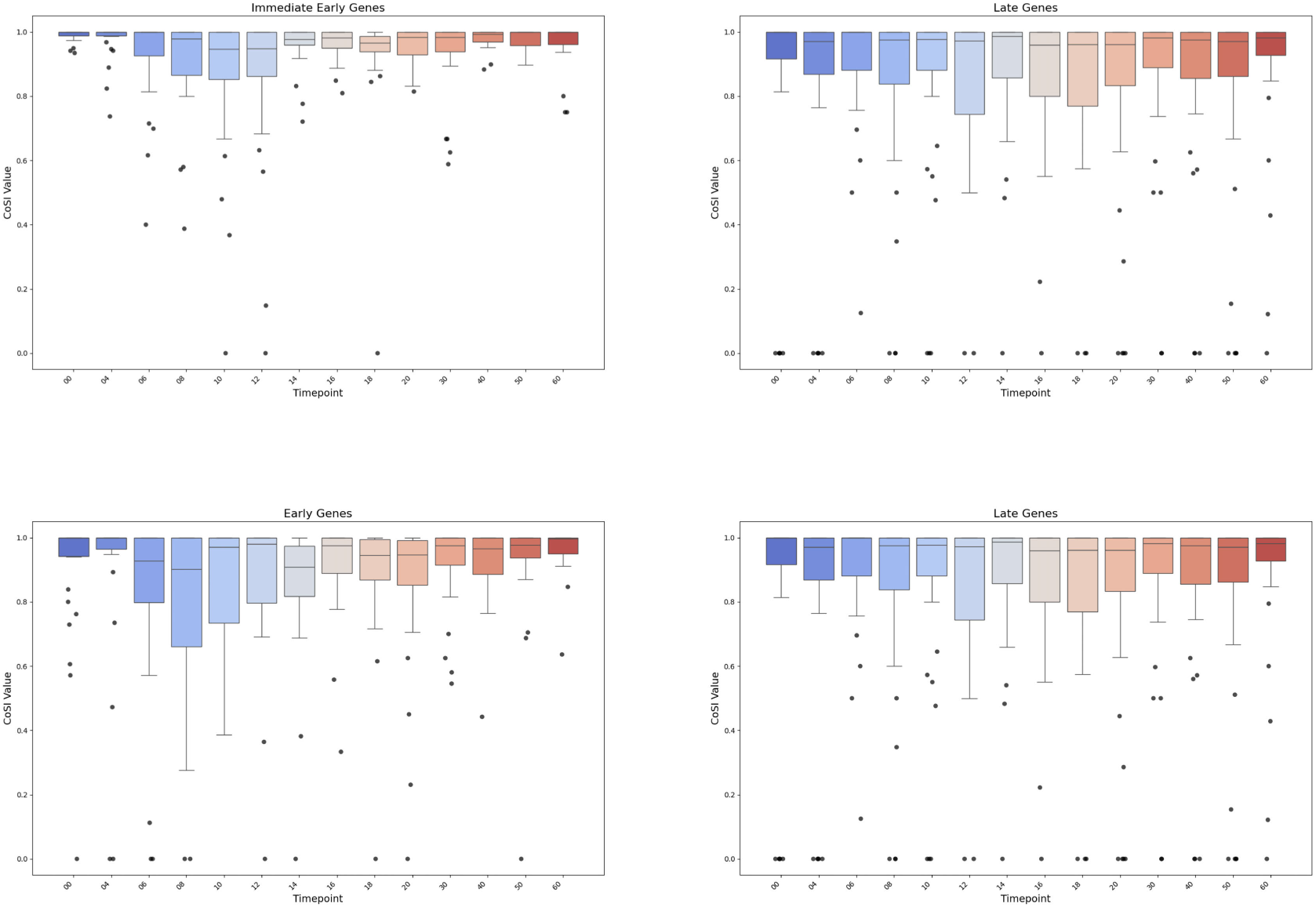
Splicing completion across kinetic gene expression groups. Box-and-whisker plots depict time-course CoSI values for introns from inflammatory gene cohorts. Each point represents the CoSI value of an individual intron at a given time point. Plots are grouped by expression category: Immediate Early (top left), Early (top right), Intermediate (bottom left), and Late (bottom right).

**Supplementary Figure 7.**
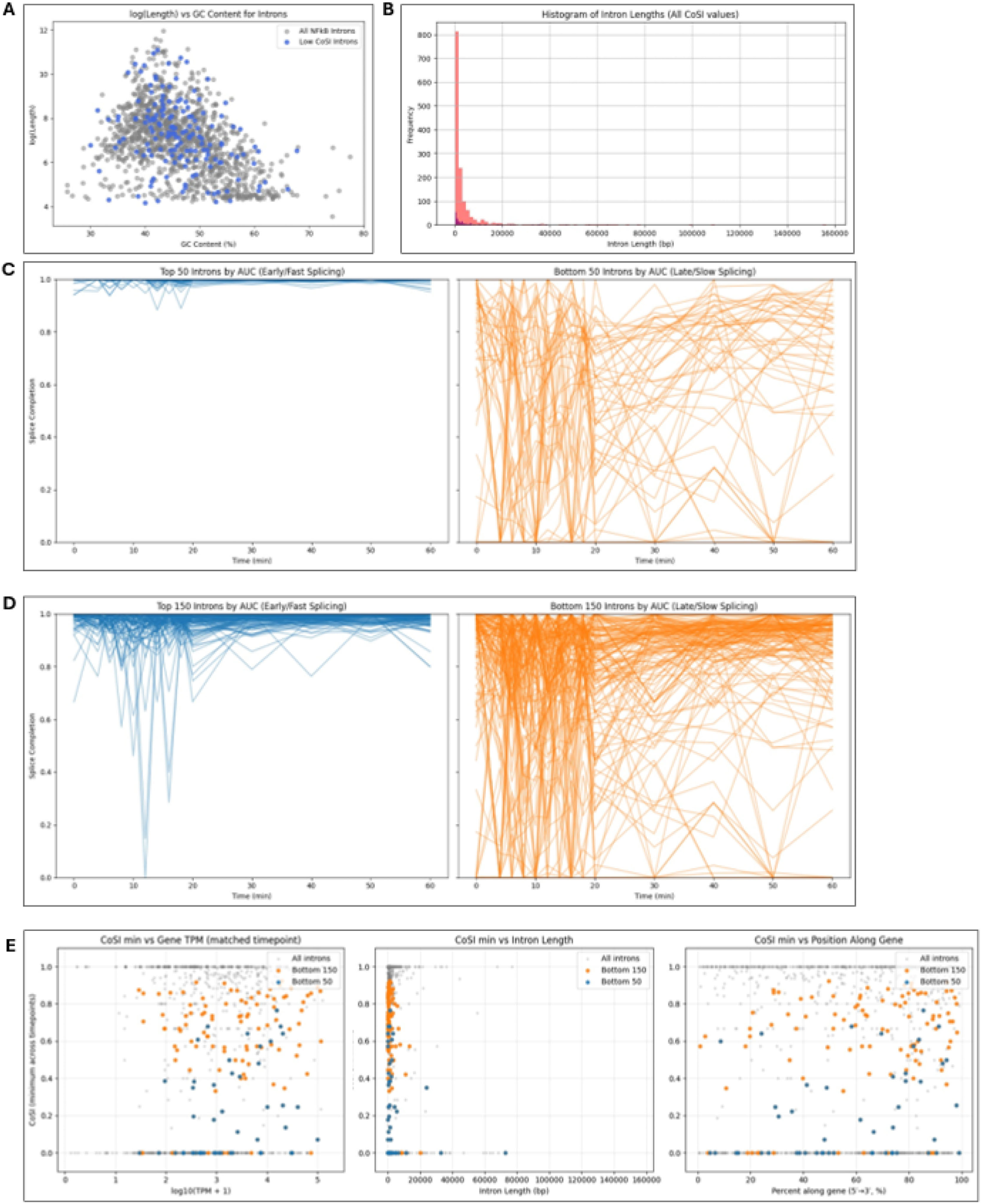
Relationships between intron length, GC content, transcript position, and splicing kinetics. (A) Scatterplot of intron length (y-axis) versus GC content (%) (x-axis) for the 150 slowest-splicing introns. Pearson *𝑟*= −0.44 (*𝑝*= 2.76×10^−45^) and Spearman *𝑟* = −0.44 (*𝑝* = 1.34×10^−46^). (B) Histogram of intron length distributions (100- nt bins) for all 1,098 introns (red) and the 150 slowest-splicing introns (blue). Length distributions are largely similar between cohorts. (C) Splice completion (CoSI) trajectories across the time course for the 50 fastest (left, blue) and 50 slowest (right, orange) introns, ranked by area under the CoSI curve (AUC). (D) Same analysis as in (C) but extended to the top 150 fastest and bottom 150 slowest introns. (E) Minimum CoSI values for each intron plotted against transcript abundance (TPM, left), intron length (middle), and position along the transcript (5^′^ → 3^′^ %, right) for the bottom 150 and bottom 50 introns. Correlations: TPM—Bottom 150: Pearson *𝑟*= 0.204 (*𝑝*= 1.24×10^−2^), Spearman *𝑟* = 0.203 (*𝑝* = 1.27×10^−2^); Bottom 50: Pearson *𝑟*= 0.276 (*𝑝*= 5.25×10^−2^), Spearman *𝑟* = 0.316 (*𝑝* = 2.53×10^−2^). Length—Bottom 150: Pearson *𝑟* = −0.242 (*𝑝* = 2.88×10^−3^), Spearman *𝑟* = −0.266 (*𝑝* = 9.92×10^−4^); Bottom 50: Pearson *𝑟* = −0.160 (*𝑝* = 0.268), Spearman *𝑟* = −0.171 (*𝑝* = 0.235). Transcript position—Bottom 150: Pearson *𝑟*= 0.260 (*𝑝*= 1.33×10^−3^), Spearman *𝑟* = 0.231 (*𝑝* = 4.44×10^−3^); Bottom 50: Pearson *𝑟*= 0.259 (*𝑝*= 6.90×10^−2^), Spearman *𝑟* = 0.318 (*𝑝* = 2.43×10^−2^).

**Supplementary Figure 8.**
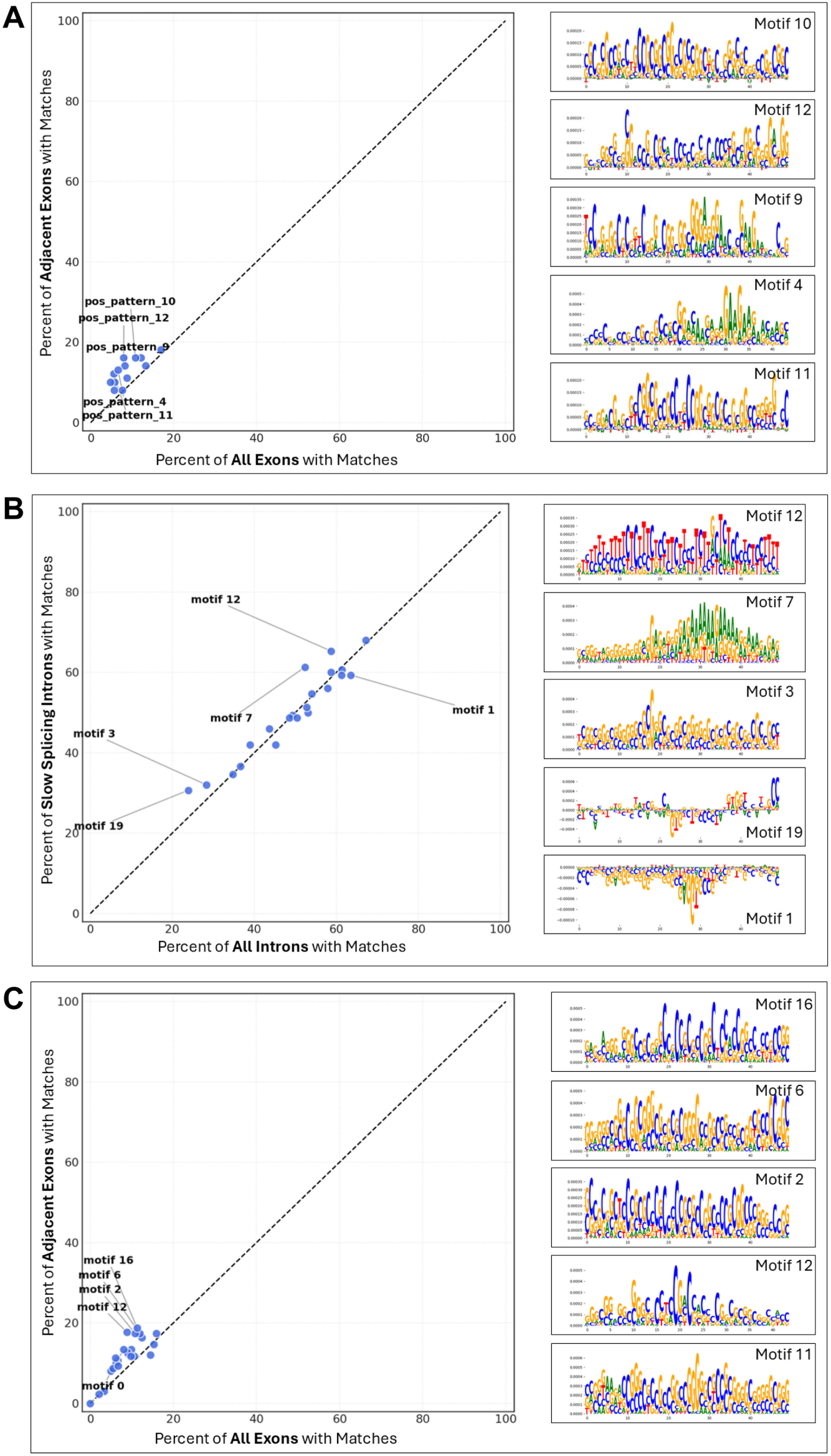
Enrichment of sequence motifs near slow-splicing introns. Scatterplots show the percent representation of position weight matrices (PWMs) scanned across slow-splicing introns compared to all introns genome-wide using FIMO. Sequence logos of the top five enriched PWMs are displayed to the right of each scatterplot. (A) Scatterplot and enriched motifs identified in exons adjacent to the 50 slowest-splicing introns. (B) Scatterplot and enriched motifs identified within the 150 slowest-splicing introns. (C) Scatterplot and enriched motifs identified in exons adjacent to the 150 slowest-splicing introns.

## Supplementary Table Legends

**Supplementary Table 1.**
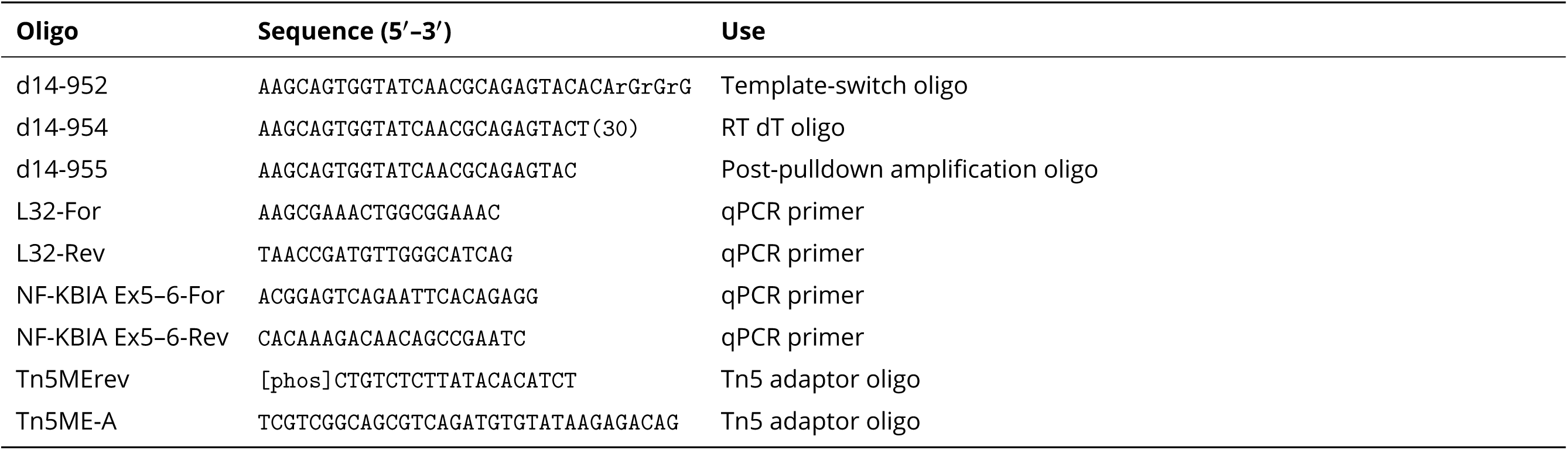
Oligonucleotide sequences used.

**Supplementary Table 2.**
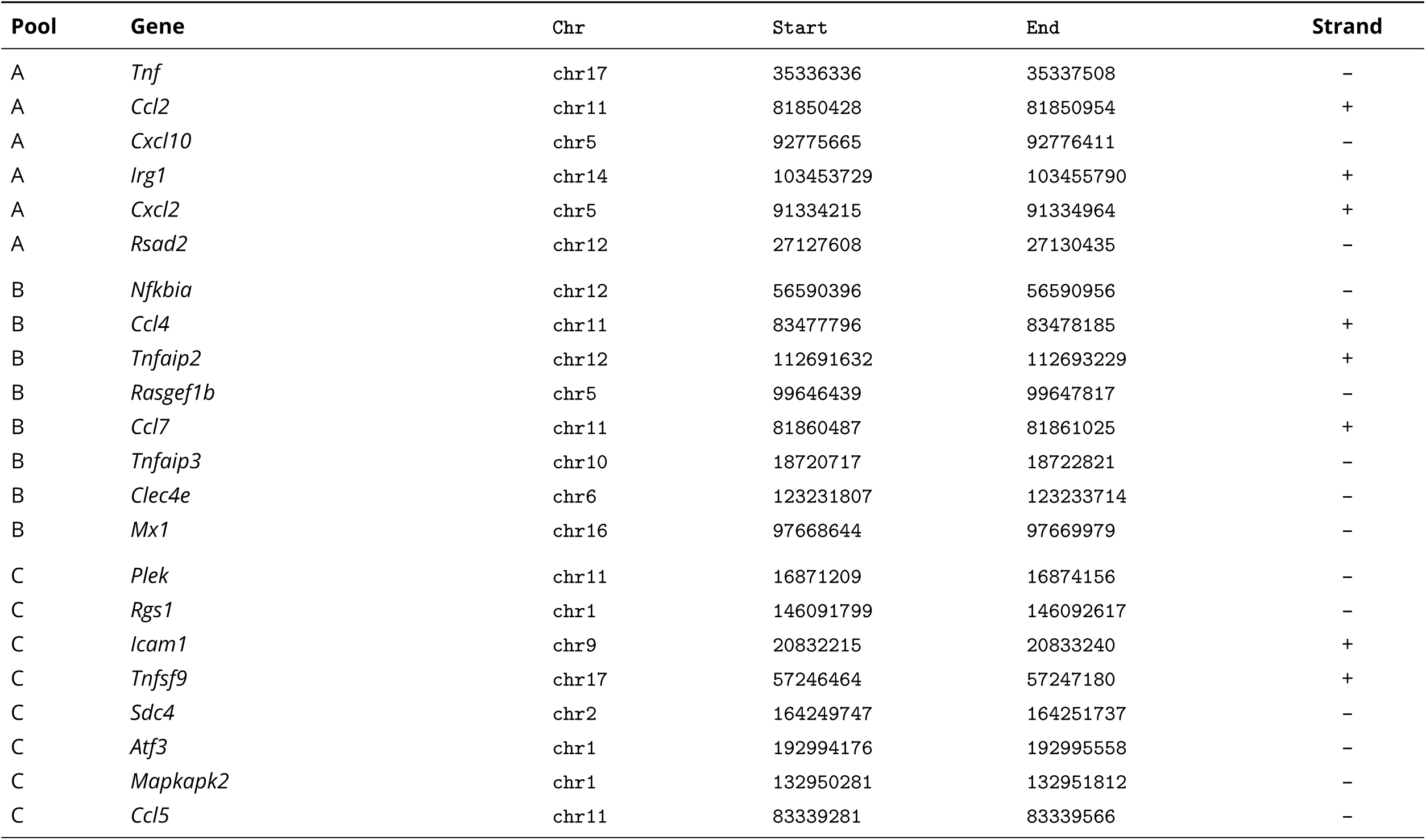
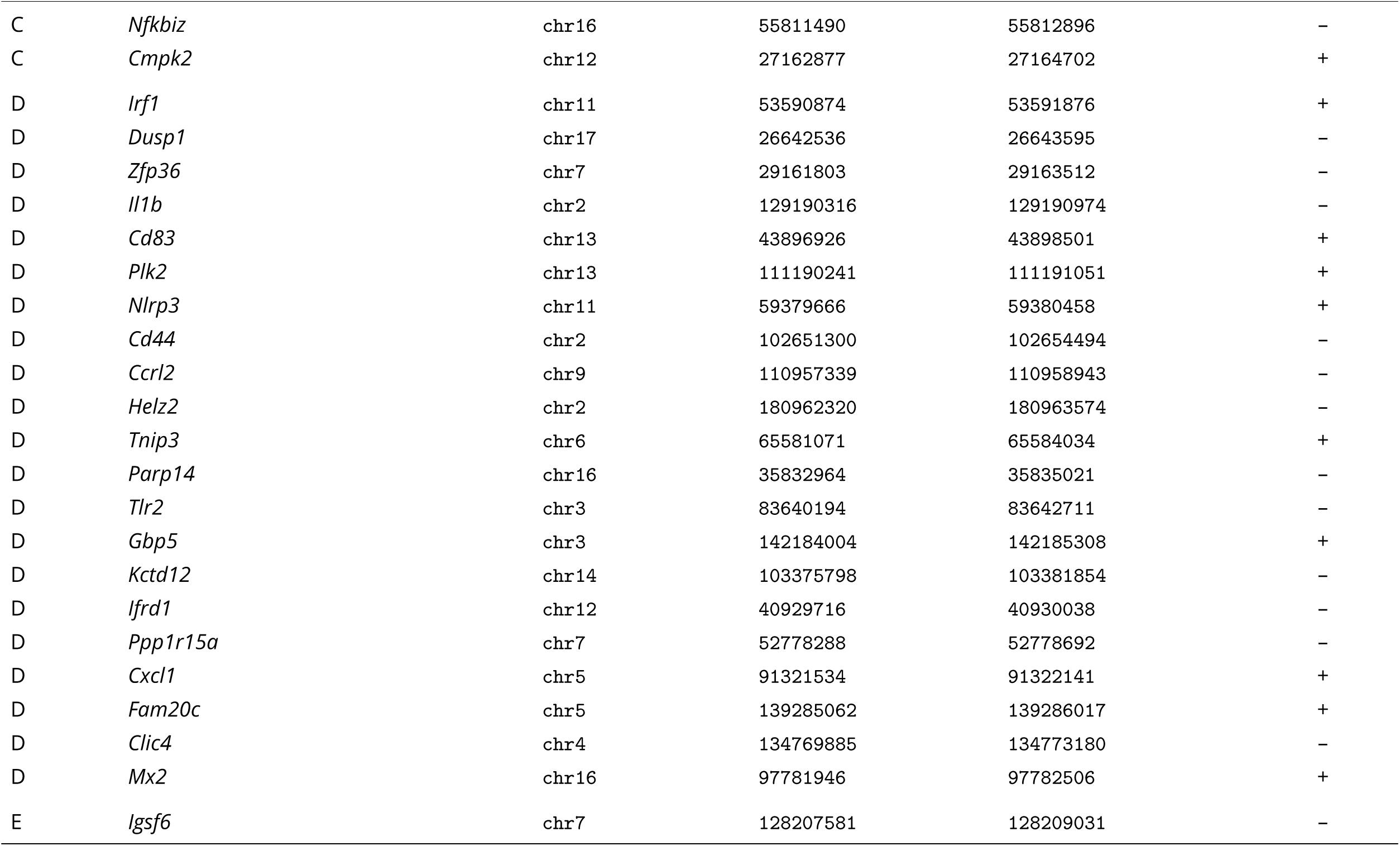
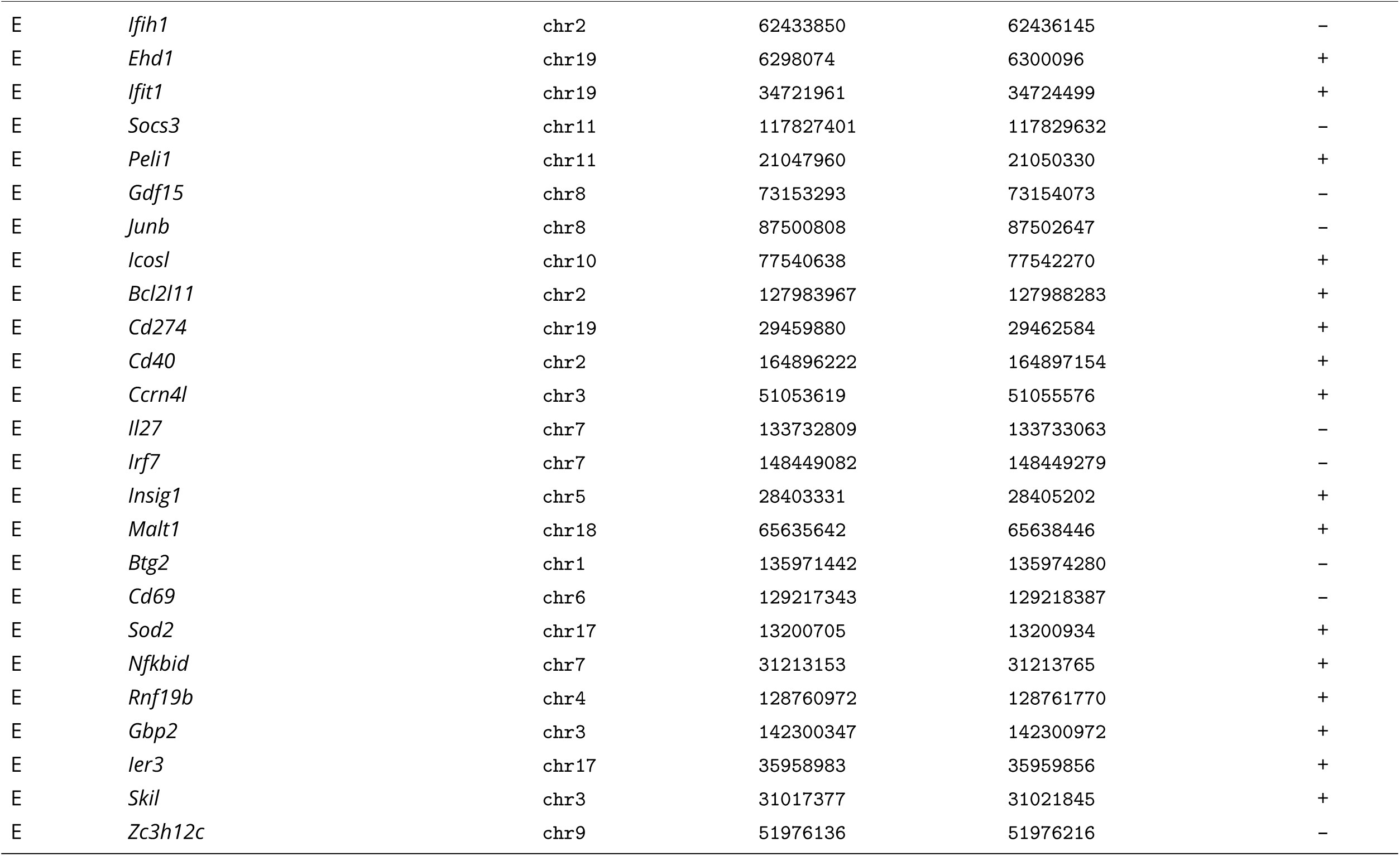
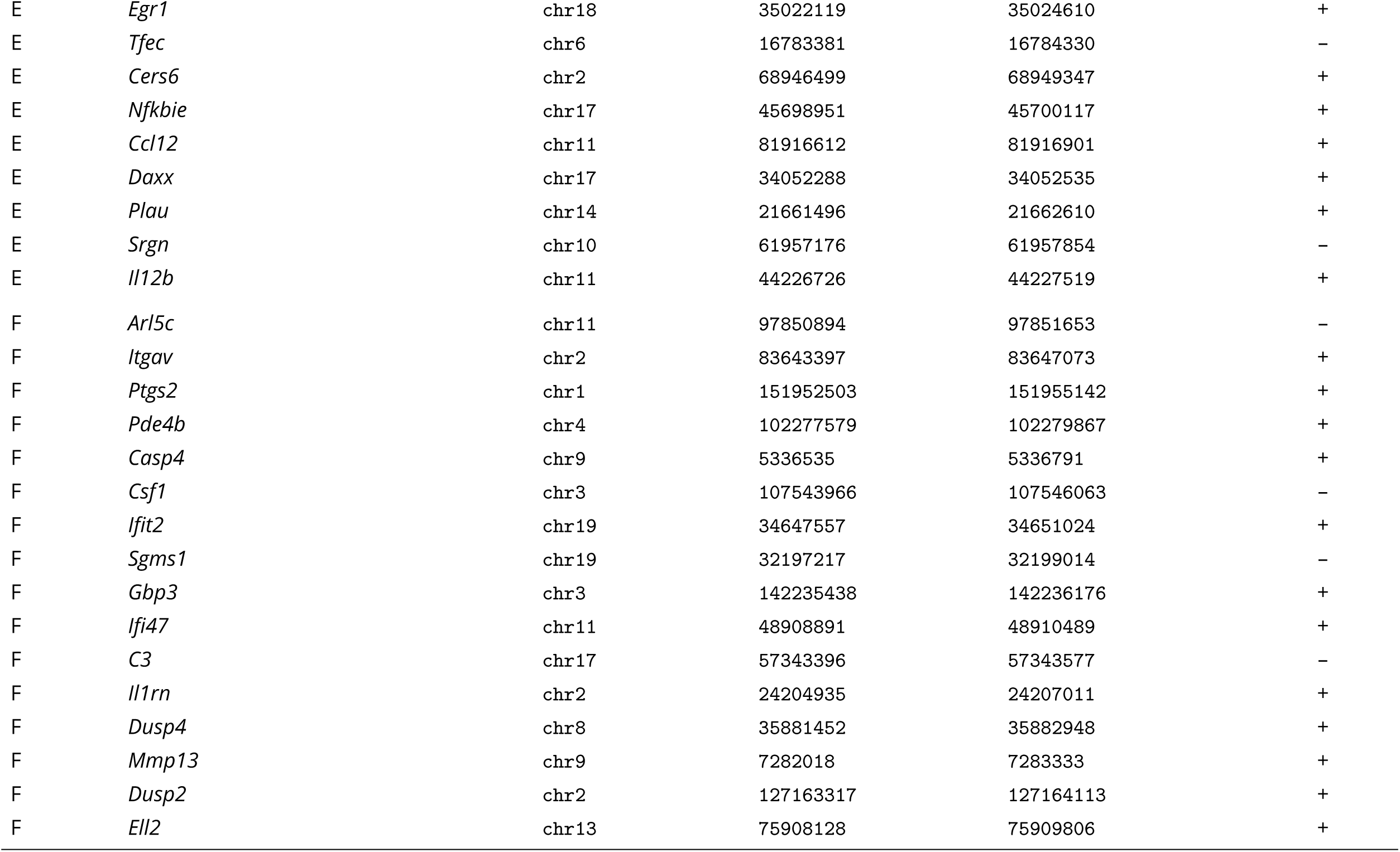
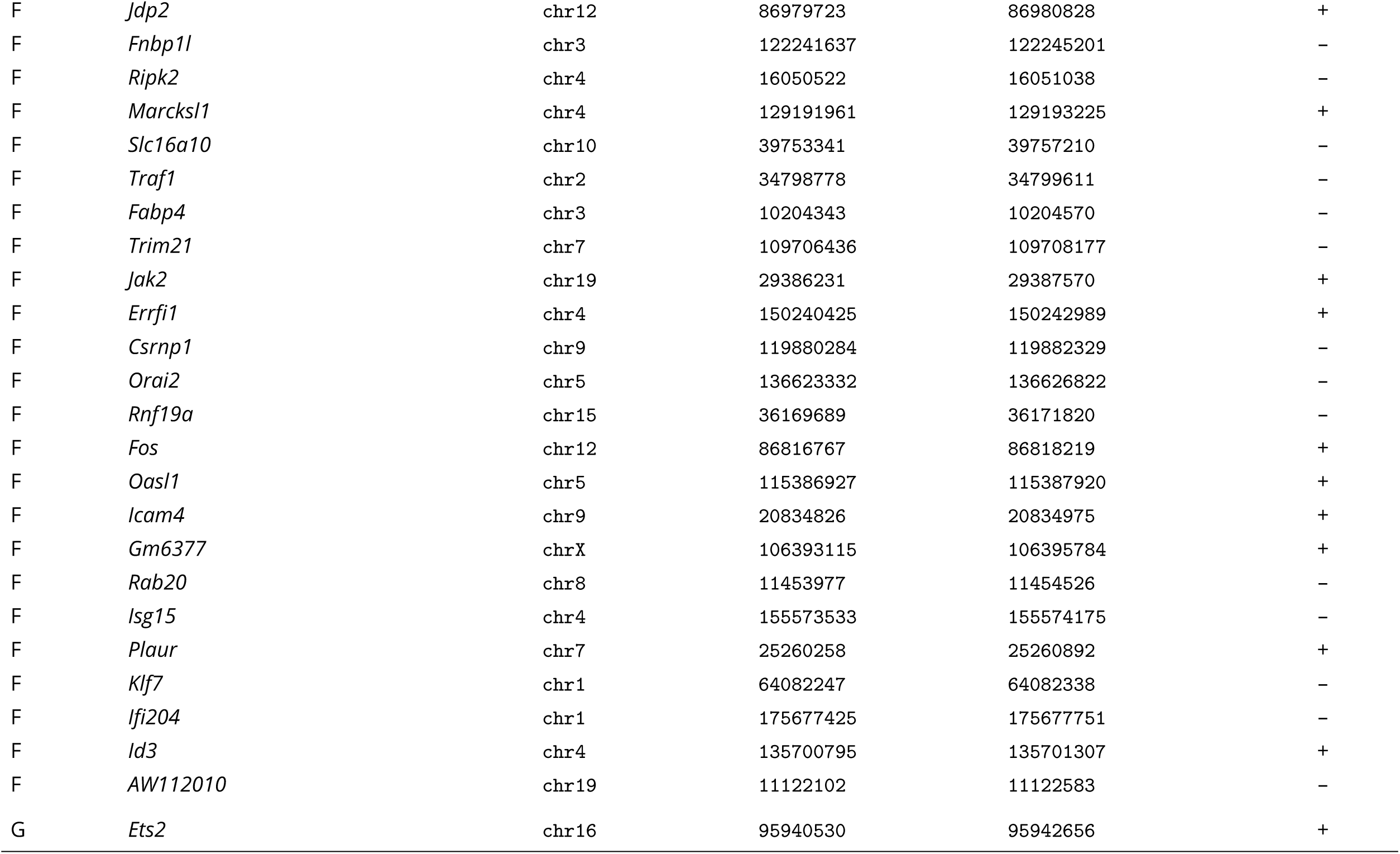
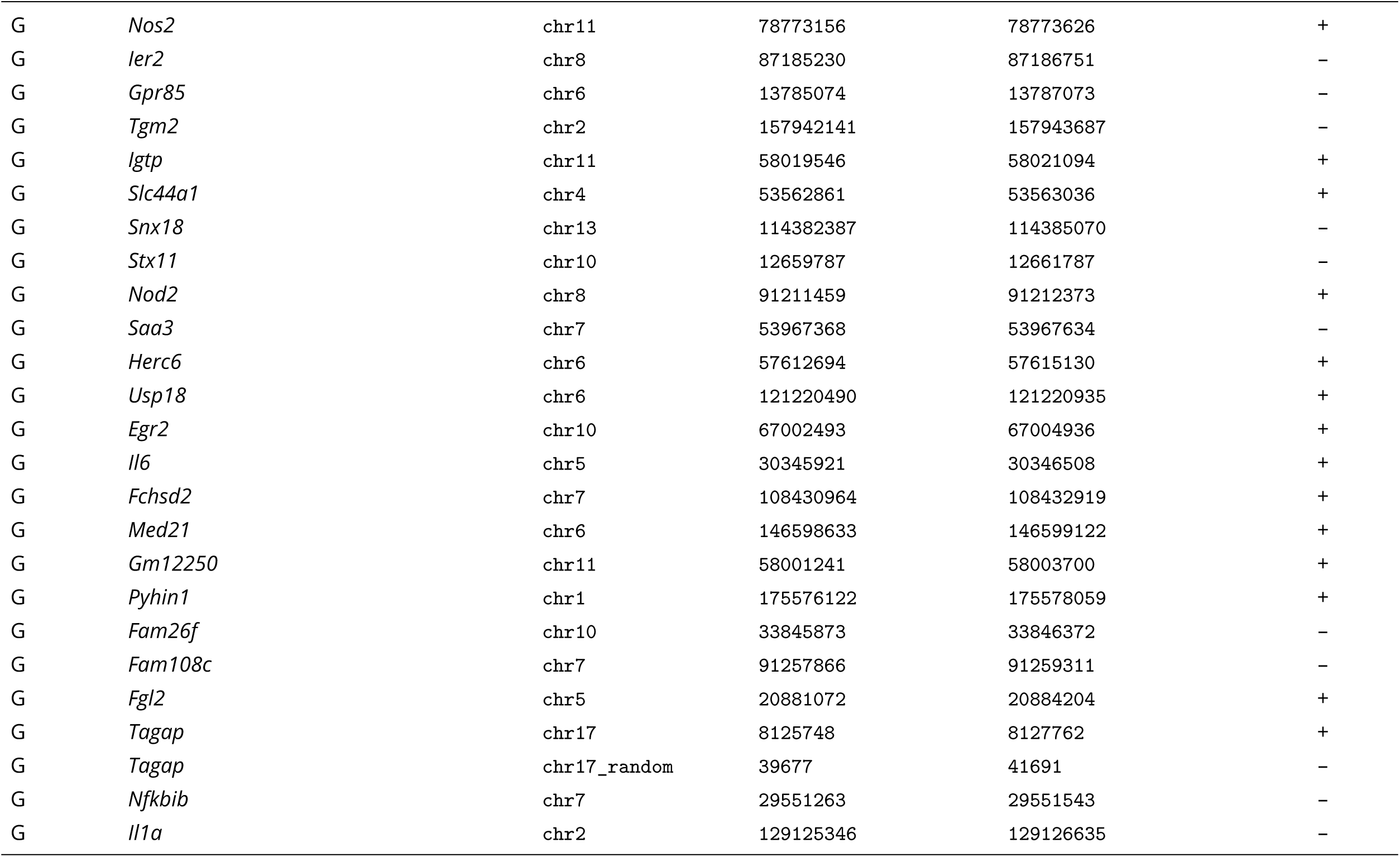
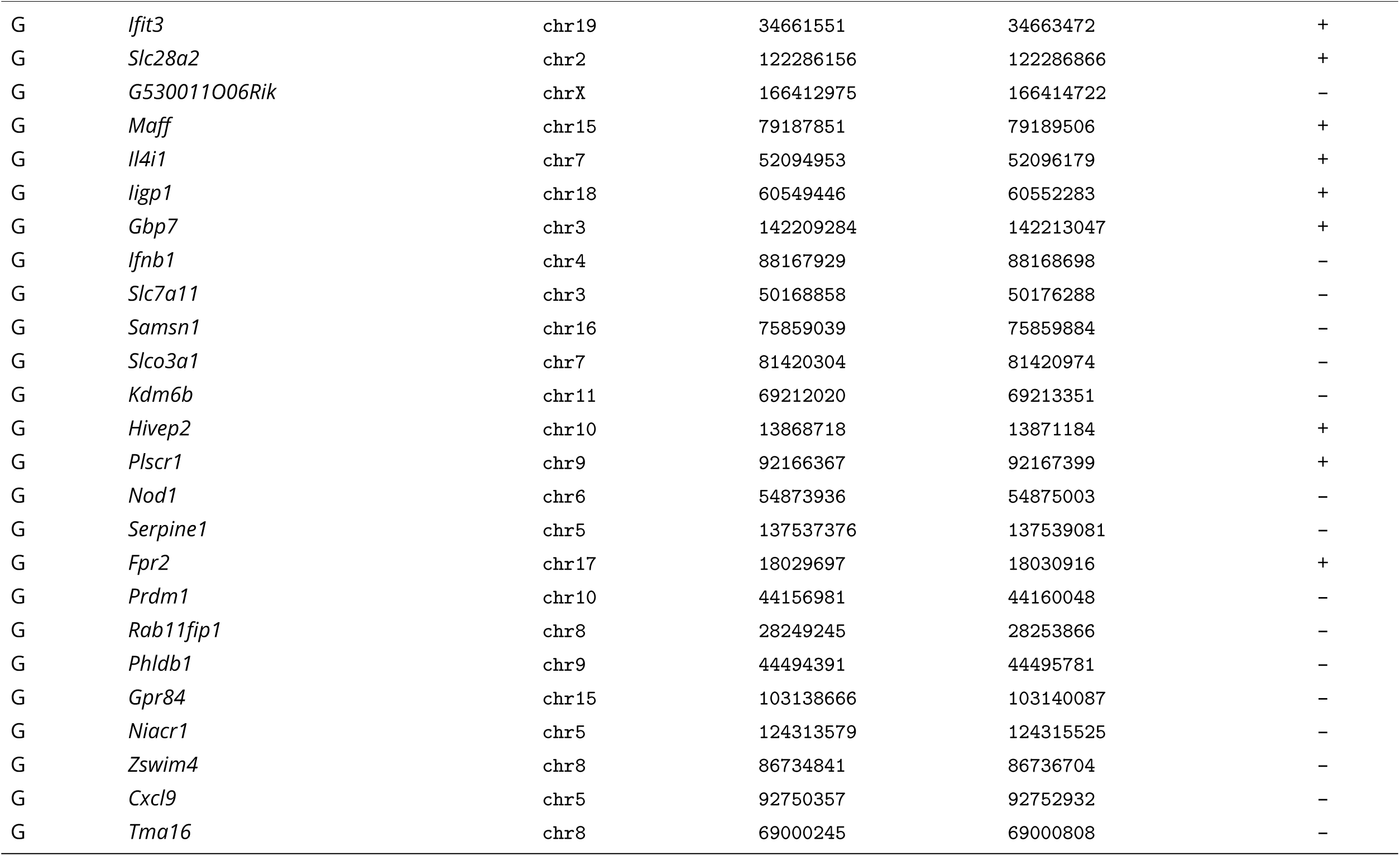
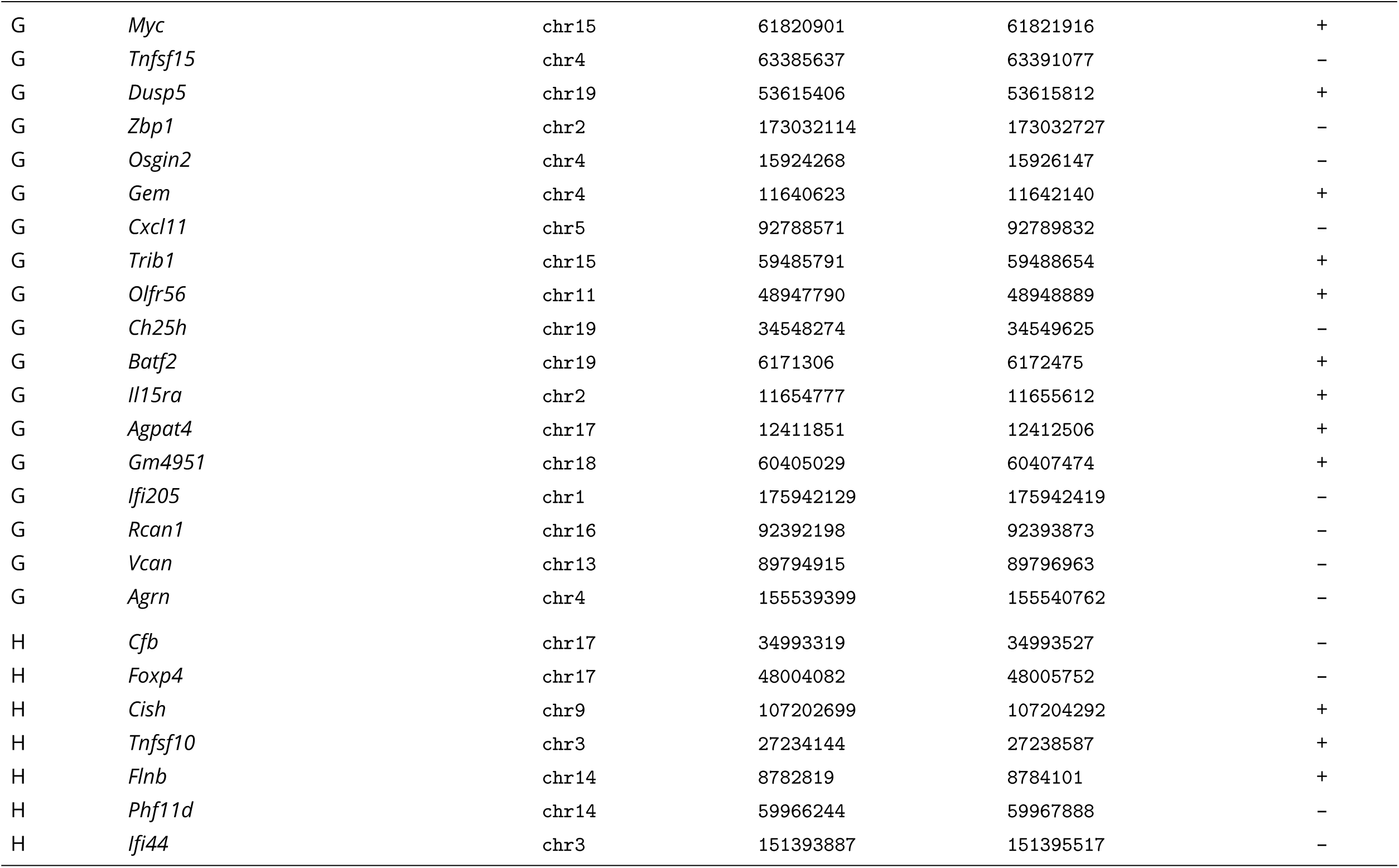
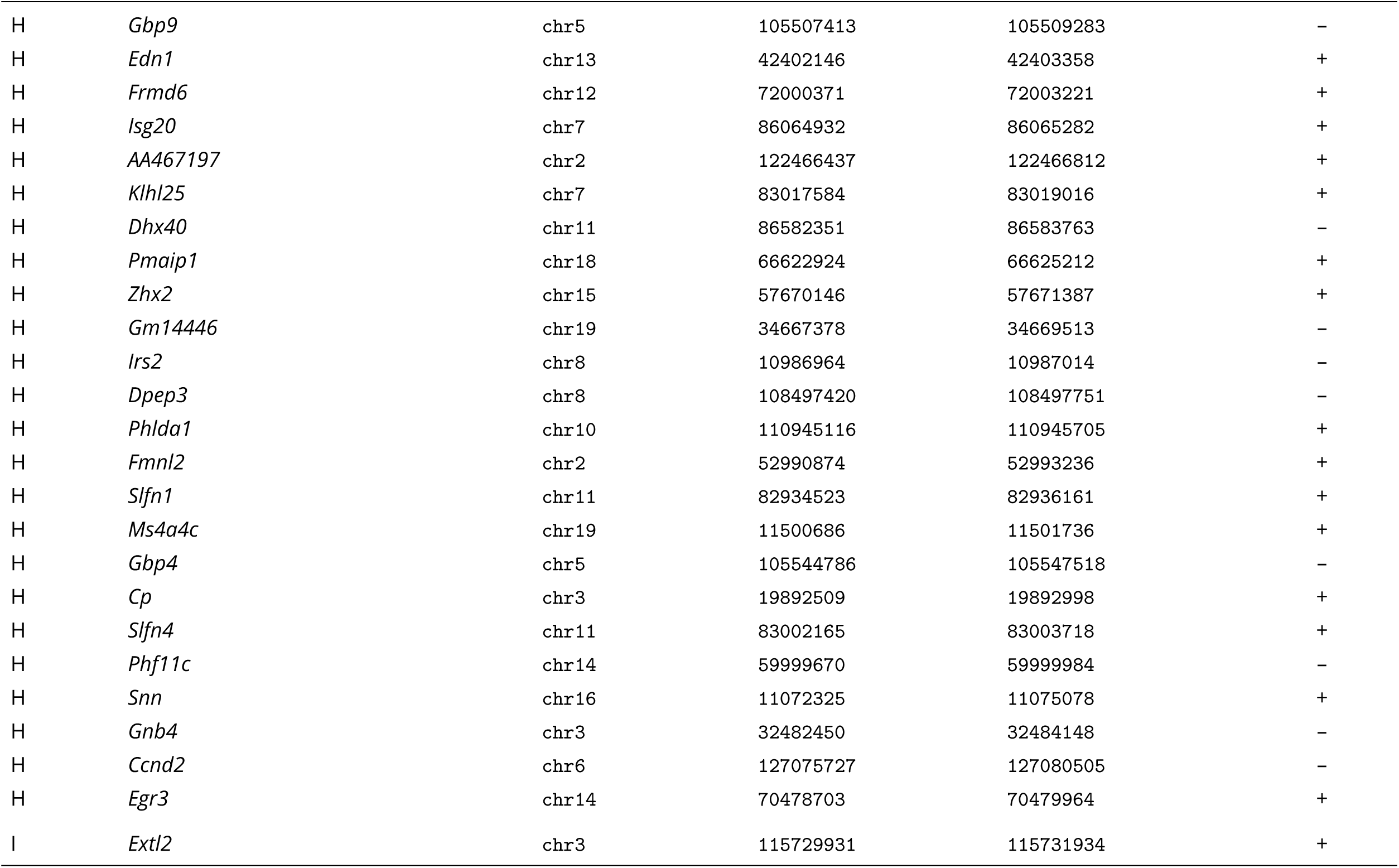
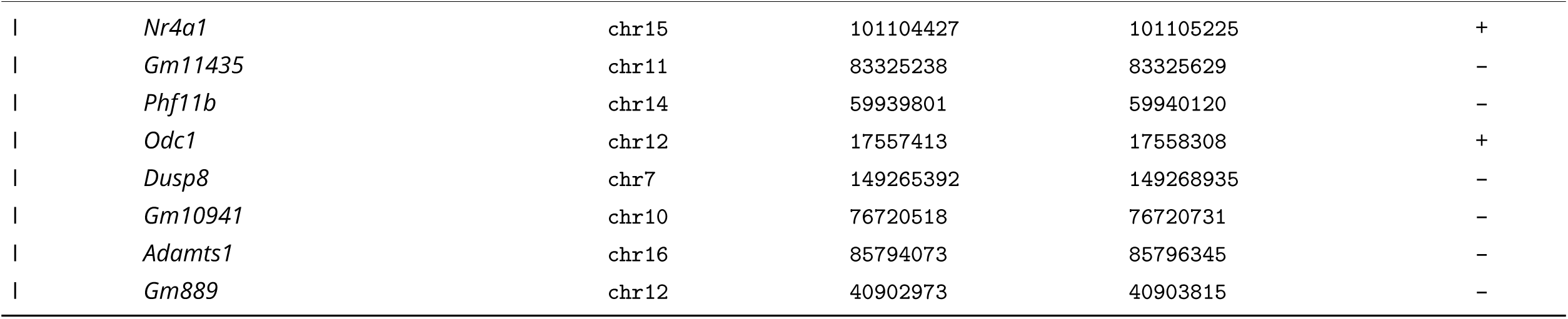
Exons of interest for hybrid capture.

